# DHX36 binding at G-rich sites in mRNA untranslated regions promotes translation

**DOI:** 10.1101/215541

**Authors:** Markus Sauer, Stefan Juranek, Hinke G. Kazemier, Daniel Benhalevy, Xiantao Wang, Markus Hafner, Katrin Paeschke

## Abstract

Translation efficiency can be affected by mRNA stability and secondary structures, including so-called G-quadruplex (G4) structures. The highly conserved and essential DEAH-box helicase DHX36/RHAU is able to resolve G4 structures on DNA and RNA *in vitro*, however a system-wide analysis of DHX36 targets and function is lacking. We globally mapped DHX36 occupancy in human cell lines and found that it preferentially binds to G-rich sequences in the coding sequences (CDS) and 5' and 3' untranslated regions (UTR) of more than 4,500 mRNAs. Functional analyses, including RNA sequencing, ribosome footprinting, and quantitative mass spectrometry revealed that DHX36 decreased target mRNA stability. However, target mRNA accumulation in DHX36 KO cells did not lead to a significant increase in ribosome footprints or protein output indicating that they were translationally incompetent. We hypothesize that DHX36 resolves G4 and other structures that interfere with efficient translation initiation.

## INTRODUCTION

All messenger RNAs (mRNAs) are coated with a dynamically changing repertoire of RNA binding proteins (RBP) to form ribonucleoprotein particles (RNP) that mediate mRNA splicing, editing, transport, turnover, and translation (Gerstberger et al., 2014; Kastelic and Landthaler, 2017; Keene, 2007). Many of these processes require energy-driven remodeling of RNP composition or RNA secondary structures by RNA helicases, a large class of motor proteins consisting of at least 95 members in humans (Gerstberger et al., 2014; Jankowsky, 2011; Umate et al., 2011). Consequently, dysregulation of helicase activity is associated with multiple diseases, including various cancer types (Fuller-Pace, 2013; Steimer and Klostermeier, 2012). For example, overexpression of eIF4A, the RNA helicase involved in translation initiation, promotes malignant transformation by increasing the protein expression of a number of oncogenes containing potentially G-quadruplex (G4) forming structures in the 5’ untranslated region (UTR) of their mRNA (Wolfe et al., 2014).

G4 structures are stable DNA or RNA secondary structures that at their core contain stacked guanine tetrads built by Hoogsteen hydrogen bonding (Bochman et al., 2012; Millevoi et al., 2012; Rhodes and Lipps, 2015). The human genome is predicted to encode over 13,000 RNA-G4 structures (Kwok et al., 2016), which may impact a wide range of processes, including mRNA 3´end processing, or telomerase activity (Millevoi et al., 2012; Rhodes and Lipps, 2015; Song et al., 2016). Most intensively studied are the effects of RNA G4 structures on translation: they are postulated to influence cap-independent translation by altering IRES (Internal Ribosome Entry Site) recognition of viral and cellular transcripts (Baird et al., 2006; Bonnal et al., 2003; Cammas et al., 2015; Morris et al., 2010), or, depending on the location of the G4 structure, they block translational elongation (Endoh and Sugimoto, 2016; Thandapani et al., 2015). These findings suggest that RNA G4 structures perform important posttranscriptional regulatory functions. Nevertheless, Guo and Bartel noted that in eukaryotes RNA G4 structures are globally unfolded *in vivo* (J. U. Guo and Bartel, 2016) and likely require a dedicated machinery to resolve them and modulate thier effects on cellular output. To this date only a handful of proteins, including the helicase DHX36, were shown to disrupt RNA G4 structures *in vitro* (Sauer and Paeschke, 2017).

DHX36 is a 3’-5’ DEAH-box helicase conserved in eukaryotes down to choanoflagellates (Lattmann et al., 2010). Loss of DHX36 in mice is embryonically lethal (Lai et al., 2012) and it is required for cardiac development (Nie et al., 2015). DHX36 is found upregulated in most breast cancer cell lines, as well as in more than 30% of human lung cancers deposited on the cBioPortal (Gao et al., 2013; Huang et al., 2012). DHX36 has two alternative gene names: RNA helicase associated with AU-rich elements (RHAU), based on its interaction with the AU-rich element of the PLAU mRNA (Tran et al., 2004), and G4 resolvase 1 (G4R1), due to its ability to unwind DNA and RNA G4 structures in HeLa cell lysate in an ATP-dependent manner (Creacy et al., 2008; Vaughn et al., 2005). In addition to its helicase core region DHX36 contains a characteristic 13 amino acid (aa) long RHAU-specific motif (RSM), which can bind RNA G4 structures *in vitro* with picomolar affinities (K_*D*_ of ~39-70 pM) (Booy et al., 2012; Chalupníková et al., 2008; Creacy et al., 2008; Lattmann et al., 2010) (Fig. 1A).

**Figure 1:**
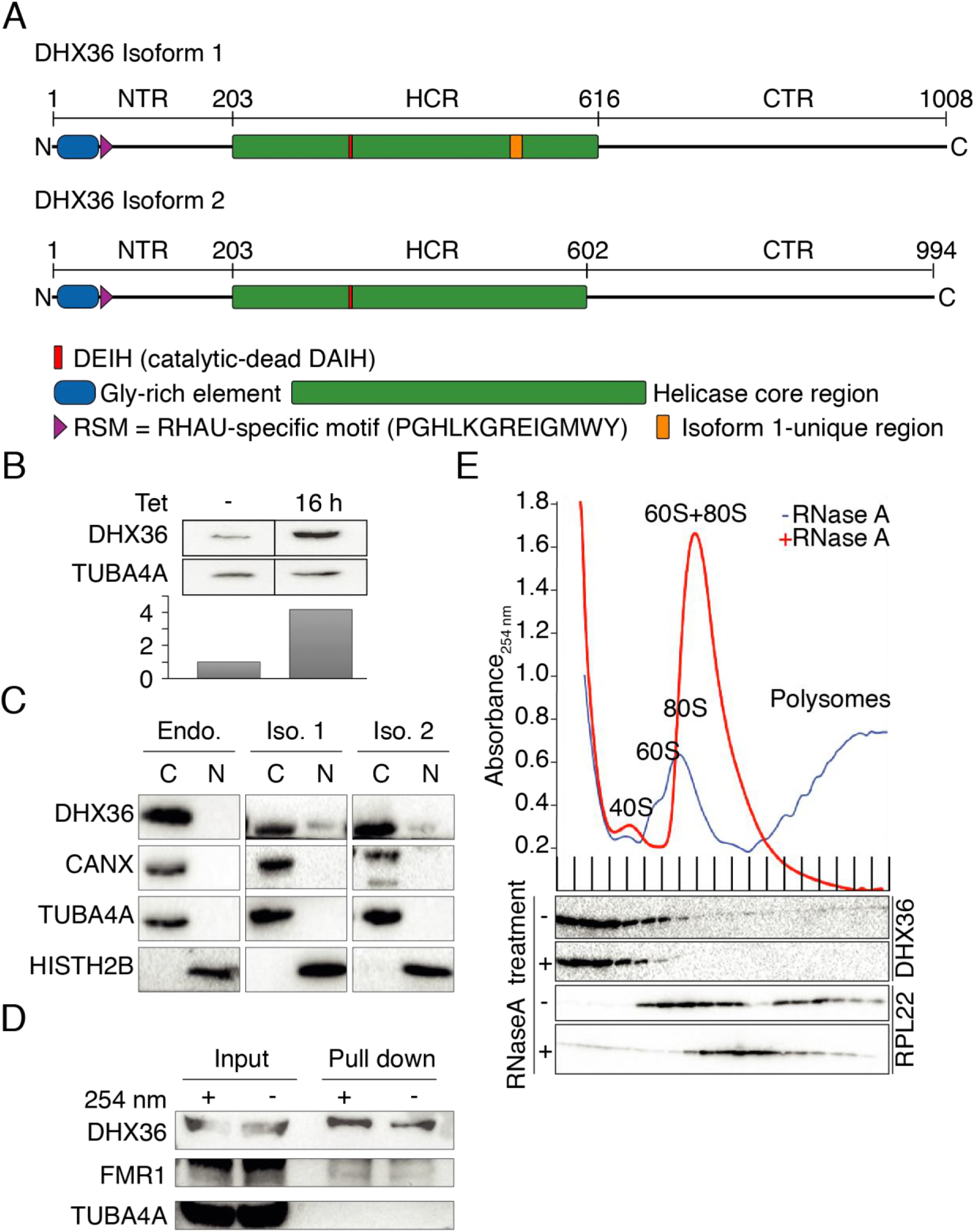
DHX36 is a mainly cytoplasmic DEAH-type helicase and does not interact with the polysome. (**A**) Schematic representation of DHX36 isoform1 and 2. The RNA binding Gly-rich element (blue) and RS motif (purple) are indicated. 14 additional aa in the helicase core region (green) of isoform 1 are marked in orange. Conserved Walker-B box (red, wildtype sequence DEIH, mutant sequence DAIH) is indicated in red. (**B**) Quantification of transgenic FLAG/HA-DHX36 expression upon 16 h induction with tetracycline. Bargraph shows DHX36 levels measured by quantitative Western blotting using an anti-DDX36 antibody and, for normalization, anti-TUBA4A antibody. (**C**) Endogenous (Endo.), as well as transgenic FLAG/HA-tagged DHX36 isoform 1 (Iso. 1) and 2 (Iso. 2) are mainly cytoplasmic in biochemical fractionation experiments from HEK293 cells. Cytosolic (C) and nuclear (N) fractions were probed with anti-HISTH2B (nuclear marker), anti-TUBA4A (cytoplasmic marker), and anti-CANX (endoplasmic reticulum marker) antibodies. Endogenous DHX36 and transgenic FH-DHX36 isoforms 1 and 2 were detected with an anti-DHX36 and anti-HA-antibodies, respectively. (**D**) DHX36 can be co-purified with polyadenylated RNA with and without UV crosslinking. The RBP FMR1 served as positive, TUBA4A as negative control, respectively. (**E**) UV absorbance at 254 nm of RNaseA-treated and untreated HEK293 cell extracts separated by sucrose gradient centrifugation are shown. Peaks of UV absorbance corresponding to 40S, 60S, 80S ribosomes, and polysomes are indicated. Western blots probed for DHX36 and RPL22 are shown below.

Most proposed molecular functions for DHX36 focus on its ability to specifically bind and unwind DNA and RNA G4 structures *in vitro* (Booy et al., 2012; Chalupníková et al., 2008; Creacy et al., 2008; Lattmann et al., 2010). Individual studies suggested that DHX36 is also able to resolve G4 structure formation *in vivo*: e.g. DHX36 was able to interact with transfected G4-forming segments of human telomerase RNA (TERC) in cultured cells and TERC mutations abolishing G4 formation *in vitro* affected cell proliferation (Booy et al., 2012; Lattmann et al., 2011; Sexton and Collins, 2011). Fusing potentially G4-forming fragments to luciferase or GFP sequences, or placing them into the promotor regions of reporter constructs affected reporter gene expression in a DHX36-dependent manner (Booy et al., 2014; Huang et al., 2012; Nie et al., 2015). In total, 27 direct RNA targets for DHX36 were previously determined (Booy et al., 2014; Nie et al., 2015; Sexton and Collins, 2011; Tran et al., 2004) (Suppl. Table S1). However, considering the large number of predicted RNA G4 structures (J. U. Guo and Bartel, 2016; Kwok et al., 2016) the full complement of DHX36 targets and its impact on gene regulation remain unknown.

Here we aimed to obtain comprehensive insight into DHX36 targets and function using biochemical and systems-wide approaches. We identified DHX36 as a predominantly cytoplasmic RBP and profiled its RNA targets on a transcriptome-wide scale at single nucleotide (nt) level resolution by Photoactivatable Ribonucleoside Enhanced Crosslinking and Immunoprecipitation (PAR-CLIP) (Hafner et al., 2010) using the wildtype protein as well as the catalytically inactive E335A mutant (Booy et al., 2014). PAR-CLIP revealed that DHX36 bound tens of thousands of G-rich sites distributed over mature mRNAs. DHX36 binding sites coincided with regions that were recently reported to form G4 and other stable structures (Kwok et al., 2016). Loss-of-function analysis with DHX36 KO cells coupled with RNA sequencing (RNA-seq), ribosome profiling (Ribo-seq), and stable isotope labeling by amino acids in cell culture (SILAC) revealed that binding of DHX36 in the 3’ and 5’ UTR resulted in higher target mRNA translational efficiency, while at the same time decreasing DHX36 target mRNA abundance. Our results underline that while G-rich regions in 3’ and 5’ UTRs can increase RNA stability, they nevertheless reduce their translation competence.

## RESULTS

### DHX36 is a cytoplasmic helicase interacting with mRNA in human cultured cells

Human DHX36 was found to encode at least two splice isoforms that differ by alternative 5’ splice site usage in exon 13 that results in the exclusion of a potential nuclear localization signal (NLS) in isoform 2 (Fig. 1A) (Tran et al., 2004). Because of the suggested nuclear as well as cytoplasmic localization for DHX36 isoforms, we investigated whether the inclusion of the putative NLS could influence subcellular localization. We generated stable HEK293 cell lines expressing FLAG/HA-tagged DHX36 (FH-DHX36) isoforms 1 and 2 under control of a tetracycline inducible promoter (Spitzer et al., 2013). Upon tetracycline induction, FH-DHX36 isoforms 1 and 2 accumulate to approx. 3 to 4-fold levels compared to endogenous DHX36 in HEK293 cells (Fig. 1B), determined by quantitative western blotting using antibodies recognizing the N-terminal region of DHX36. Subcellular fractionation revealed that both transgenic DHX36 isoforms, as well as endogenous DHX36 predominantly localize to the cytoplasm (Fig. 1C).

While DHX36 has been described as both a DNA and RNA helicase *in vitro*, the major source of nucleic acids in the cytosol is RNA. Thus, we tested whether DHX36 indeed interacted with RNA, in particular mRNAs, in human cell lines. Cells were crosslinked using 254 nm UV-light in HEK293 cells followed by purification of polyadenylated RNA (Fig. 1D). We found that DHX36 was abundantly interacting with polyadenylated RNA, indicating that cytosolic mRNAs are the main targets of DHX36.

### DHX36 is found in the soluble cytoplasm and associated with the translational machinery

The interaction of DHX36 and mRNAs in the cytosol suggested a posttranscriptional regulatory function, which is in agreement with a previously proposed function of DHX36 in translational regulation (Thandapani et al., 2015). We fractionated HEK293 cells extract by ultracentrifugation through sucrose gradient to investigate whether DHX36 co-migrates with the translational machinery. In proliferating HEK293 cells approximately 90% of endogenous DHX36 migrated with the soluble cytoplasm, and the remaining 10% migrated with the monosomal and polysomal fractions (Fig. 1E). Treatment of the lysate with RNase A resulted in the collapse of polysomes and the loss of DHX36 from heavier fractions (Fig. 1E). Taken together, this data indicated that only a fraction of DHX36 is loaded on actively translated mRNAs, implying that a putative effect on translation would likely not occur at the translational elongation step.

### DHX36 binds thousands of sites on mature mRNAs

In order to reliably capture DHX36 binding sites and characterize its RNA recognition elements (RREs) we mapped the RNA interactome of DHX36 in living cells on a transcriptome-wide scale at nucleotide-resolution using 4-thiouridine (4SU) PAR-CLIP (Hafner et al., 2010). UV-crosslinking of active helicases that translocate on their RNA targets may result in the recovery of transient interactions between RNA and protein, which could complicate the determination of the preferred interaction sites. To overcome this difficulty, we performed PAR-CLIP in two stable HEK293 cell lines, either inducibly expressing FLAG-HA tagged DHX36 or the catalytically dead DHX36 E335A mutant, which is expected to remain bound at the sites of DHX36 recruitment to the mRNA.

Autoradiography of the crosslinked, ribonuclease-treated, and radiolabeled FLAG-immunoprecipitate confirmed the isolation of one main RNP depicted by a single major band at the expected size of ~130 kDa corresponding to the DHX36 and FH-DHX36 E335A RNPs (Fig. 2A). We recovered the bound RNA fragments from the FH-DHX36 and FH-DHX36 E335A RNPs of two biological replicates and generated small RNA cDNA libraries for next-generation sequencing. We used the PARalyzer software (Corcoran et al., 2011) to determine clusters of overlapping reads that contained T-to-C mutations diagnostic of the crosslinking event at higher frequencies than expected by chance (see Suppl. Table S1 for summary statistics). The biological replicates from the FH-DHX36 and FH-DHX36 E335A PAR-CLIP exhibited high correlation, with an R^2^ of 0.79 and 0.93, respectively (Suppl. Fig. S1A, B). For the FH-DHX36 and FH-DHX36 E335A PAR-CLIPs we defined as high-confidence binding sites 19,585 and 67,660 clusters, respectively, that were found in both replicates (Suppl. Table S2). Overall the binding profiles of FH-DHX36 and FH-DHX36 E335A were similar, with an R^2^ of 0.66, indicating that inactivating the helicase domain did not interfere dramatically with the binding pattern of the protein (Fig. 2B).

**Figure 2:**
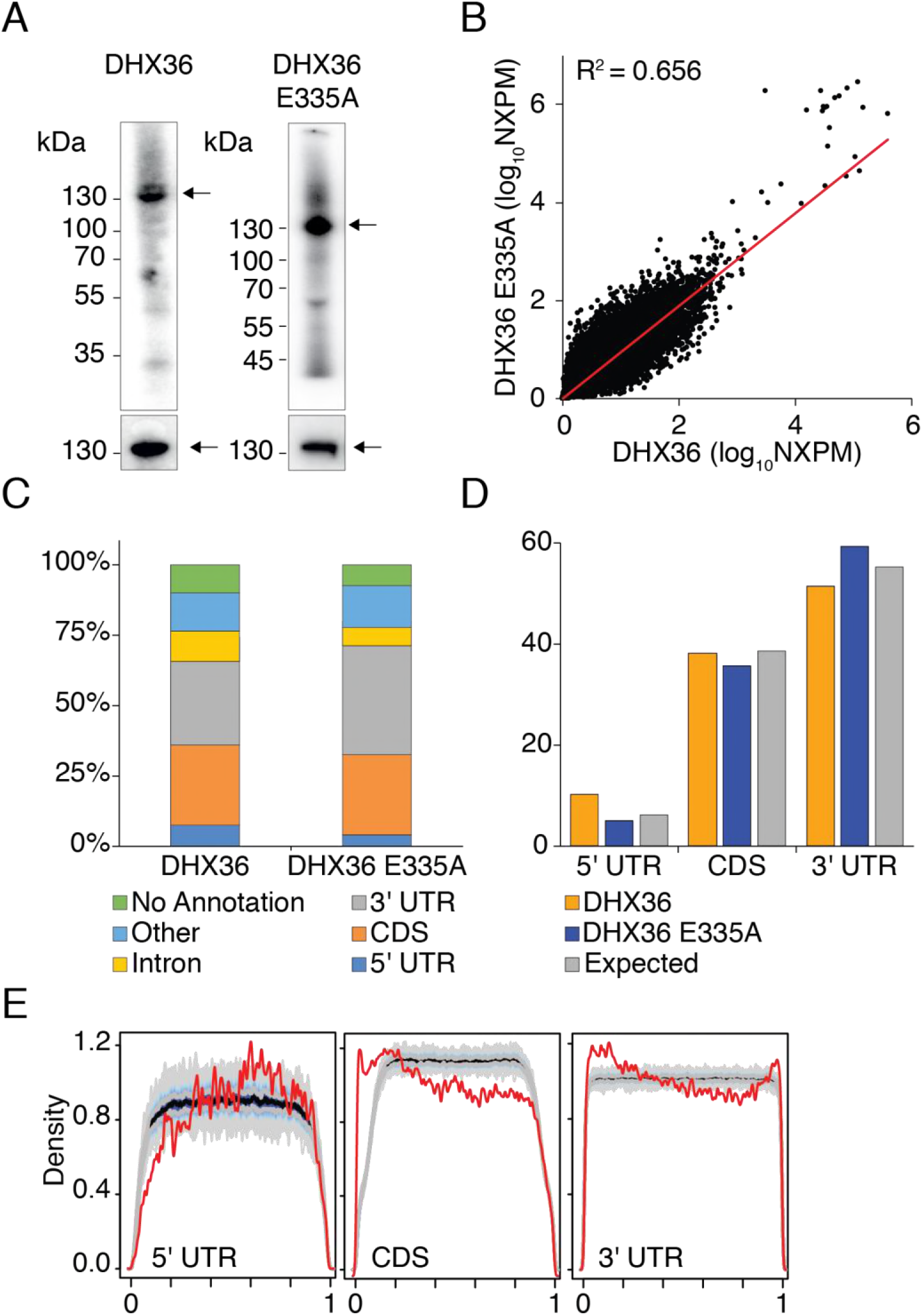
DHX36 interacts with mature mRNAs at thousands of sites. (**A**) Autoradiographs showing *in vivo* crosslinked DHX36- and DHX36 E335A-RNA RNPs from stable HEK293 cells inducibly expressing FH-DHX36 and FH-DHX36 E335A, respectevely. Black arrows indicate crosslinked DHX36-and DHX36 E335A-RNPs, respectively. Western blot analysis of immunoprecipitated FH- DHX36 and FH-DHX36 E335A is shown in the lower panel. (**B**) Scatterplot of normalized crosslinked reads per million (NXPM) from DHX36 and DHX36 E335A PAR-CLIP experiments shows high-degree of correlation of high-confidence binding sites. Correlation coefficient (R^2^) is indicated. (**C**) Distribution of PAR-CLIP-derived binding sites across different annotation categories shows that DHX36 mainly bound mature mRNA in CDS and UTR. (**D**) The distribution of DHX36 (orange) and DHX36 E335A (blue) binding sites across CDS, 3', and 5’ UTR matches the distribution expected based on the length of the annotation categories (grey). (**E**) DHX36 binds preferentially close to the start and the stop codon based on a metagene analysis of the distribution of DHX36 binding clusters on mRNAs subdivided into 5’ UTR, CDS, and 3’ UTR (red lines). The distribution of 1,000 mismatched randomized controls is shown in grey lines

Our approach succeeded to capture 22 of 27 previously published targets of DHX36, including TERC and PLAU, confirming the validity of our approach (Suppl. Table S1). Consistent with their mainly cytoplasmic localization, 70% and 73% of FH-DHX36 and FH-DHX36 E335A binding sites, respectively, mapped to the exonic regions of more than 4,500 protein-coding genes with the rest found on intronic sequences or non-coding RNAs (Fig. 2C). FH-DHX36 and FH-DHX36 E335A did not exhibit preference for the coding sequence or untranslated regions (3’ and 5’ UTR) of mRNA targets (Fig. 2D), which may be the result of FH-DHX36 interacting with RNA independent of the translational machinery. A metagene analysis revealed an enrichment of FH-DHX36 and FH-DHX36 E335A binding sites within the first 100 nucleotides of the start codon in the CDS, similar to another cytoplasmic G-rich element binding protein (Benhalevy et al., 2017), as well as directly downstream of the stop codon (Fig. 2E and Suppl. Fig. S1E).

### DHX36 binds G-rich sequences *in vivo* and can unwind G4 structures *in vitro*

Binding of FH-DHX36 and FH-DHX36 E335A to mRNAs showed no correlation to transcript length or abundance as determined by RNA-seq in HEK293 cells (Suppl. Fig. S1C, D). This suggested sequence- or structure-dependent determinants of FH-DHX36 binding, rather than unspecific interactions. Therefore, we aimed to define the preferred RRE of FH-DHX36 and FH-DHX36 E335A. We first counted the occurrence of all possible 5-mer sequences in our high-confidence binding sites and calculated their Z-score over a background of shuffled sequences of the same nucleotide composition. 5-mers that contained at least three guanines were enriched in both, FH-DHX36 and FH-DHX36 E335A, PAR-CLIPs (Fig. 3A, B and Suppl. Fig. S2A and Table S2) and in the FH-DHX36 E335A PAR-CLIP we additionally found the enrichment of 5-mers that were A/U-rich (Fig. 3C and Suppl. Table S2). We also used MEME (Bailey et al., 2009) as an alternative *in silico* approach to define the RRE. Here, the most significantly enriched RRE was also G-rich, and matched the motif for G4 formation (Todd et al., 2005) (Fig. 3D). Indeed, circular dichroism (CD) spectroscopy analysis confirmed that synthetic oligonucleotide corresponding to an representative PAR-CLIP RRE showed the characteristic spectrum of a parallel G4 structure, while sequence mutations exchanging G residues with A or C resulted in a loss of the G4 signature (Suppl. Fig. S2B). Microscale thermophoresis experiments confirmed that FH-DHX36 specifically bound the G4 forming oligonucleotides, but not the mutant s (Fig. 3E).

**Figure 3:**
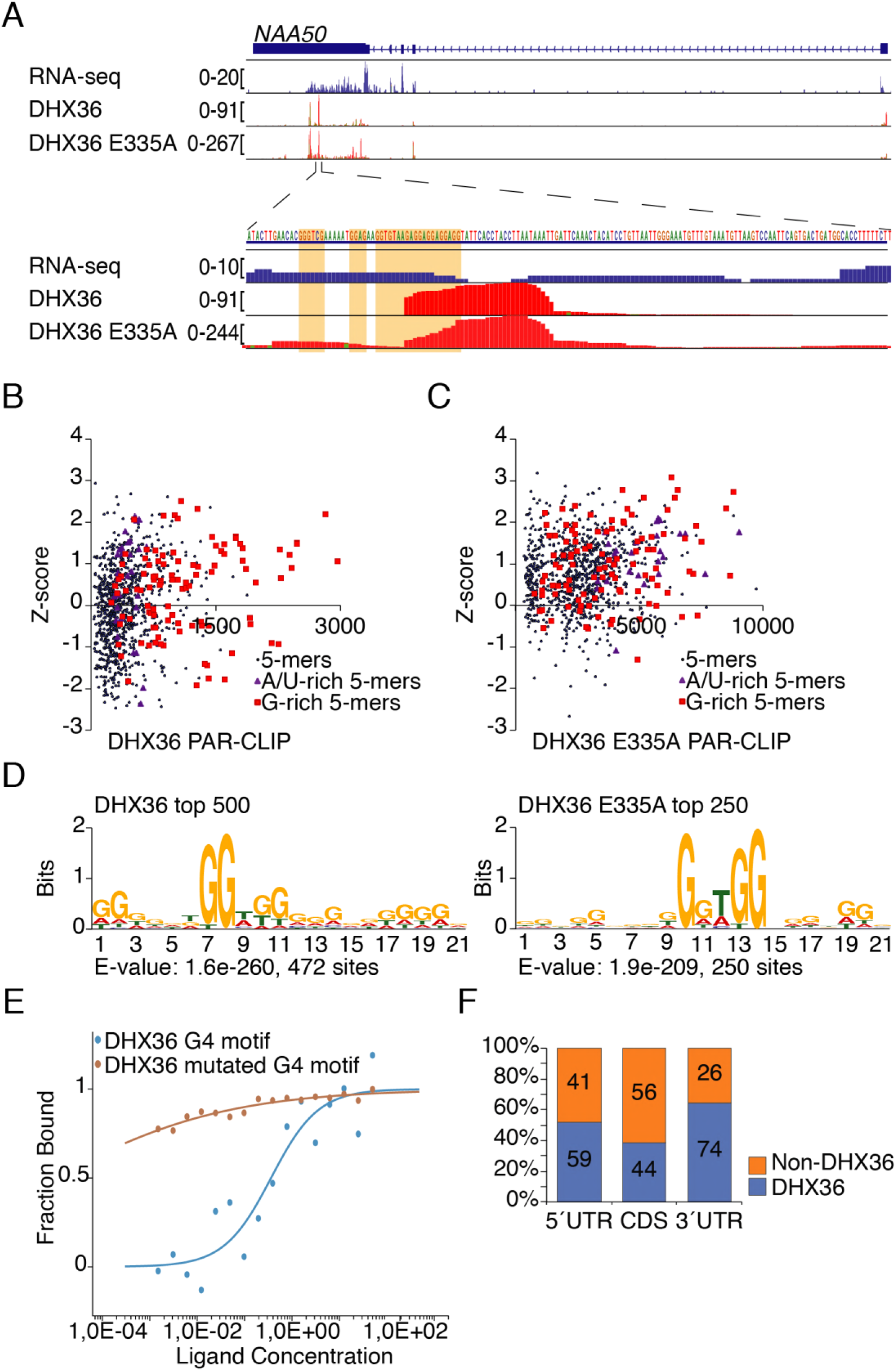
DHX36 recognizes G-rich sequence stretches on mRNA. (**A**) Top panel: Screenshot of the DHX36 and DHX36 E335A PAR-CLIP binding sites for the representative target gene NAA50. The gene structure is shown, as well as coverage from a HEK293 RNA-seq experiment. The bottom two tracks show the alignment of sequence reads with characteristic T-to-C mutations from a DHX36 and DHX36 E335A PAR-CLIP experiment. Bottom panel: Close-up of the indicated 150 nt region in the 3’ UTR of NAA50. The G-rich DHX36 binding element is highlighted in orange. **(B)** A comparison of Z-scores and occurrence of all possible 5-mers shows an enrichment of G-rich sequences in DHX36 PAR-CLIP binding sites. 5-mers containing at least three Gs (red squares) or consist of A and U- (purple triangles) are highlighted. (**C**) Same plot as in (B) for DHX36 E335A PAR-CLIP shows additional enrichment for A/U-rich 5-mers. (**D**) Weblogo of RNA recognition element of DHX36 (left) and DHX36 E335A (right) PAR-CLIP binding sites generated by MEME (p-value less than 0.0001). (**E**) Microscale thermophoresis analysis shows binding of DHX36 to the DHX36-binding motif (Cy5-AAAAAGGAGGAGGAGGAGG) but not to the mutated motif (Cy5-AAAAAGCAGCAGGAGCAGCA). (**F**) Percent of sites in the human transcriptome forming G4 structures *in vitro* categorized by 5’ UTR, CDS, and 3’ UTR (Kwok et al., 2016) found in DHX36 PAR-CLIP binding sites (blue).

Considering that this representative RRE was able to form a G4 structure we asked whether the FH-DHX36 PAR-CLIP binding sites were contained in a previously reported dataset that mapped G4 structures *in vitro* in polyadenylated RNA from HeLa cells in a transcriptome-wide manner (Kwok et al., 2016). Interestingly, 74% of the potential G4 sites found by Kwok et al. in the 3’ UTR were recovered in the FH-DHX36 E335A PAR-CLIP, despite that we were using a cell line with a different transcriptome profile for our experiments (Fig. 3F). 59% and 44% of the G4 sites in 5´UTR and CDS, respectively, also overlapped with FH-DHX36 E335A PAR-CLIP binding sites. Collectively, our *in vivo* and *in vitro* data showed that FH-DHX36 is preferentially binding G-rich elements in the CDS and UTRs which share the potential to fold into secondary RNA structures like G4 structures.

### DHX36 impacts target RNA abundance

In order to investigate gene regulatory roles of DHX36 in loss-of-function studies, we generated DHX36 knockout (KO) HEK293 cells using Cas9-mediated gene editing. Sequencing of the DHX36 locus and western blotting of the clone used in follow-up experiments confirmed an extensive deletion and loss of detectable protein (Fig. 4A, Suppl. Fig. S3A). In contrast to DHX36 transgene induction, DHX36 loss had a profound effect on the growth rate and morphology of HEK293 cells. KO cells proliferated at ~50% growth rate (Fig. 4B) and cells appeared incapable of spreading evenly in the culture dish (Suppl. Fig. S3B) consistent with a cell proliferation defect found in hematopoietic cells from conditional DHX36 KO mice (Lai et al., 2012).

**Figure 4:**
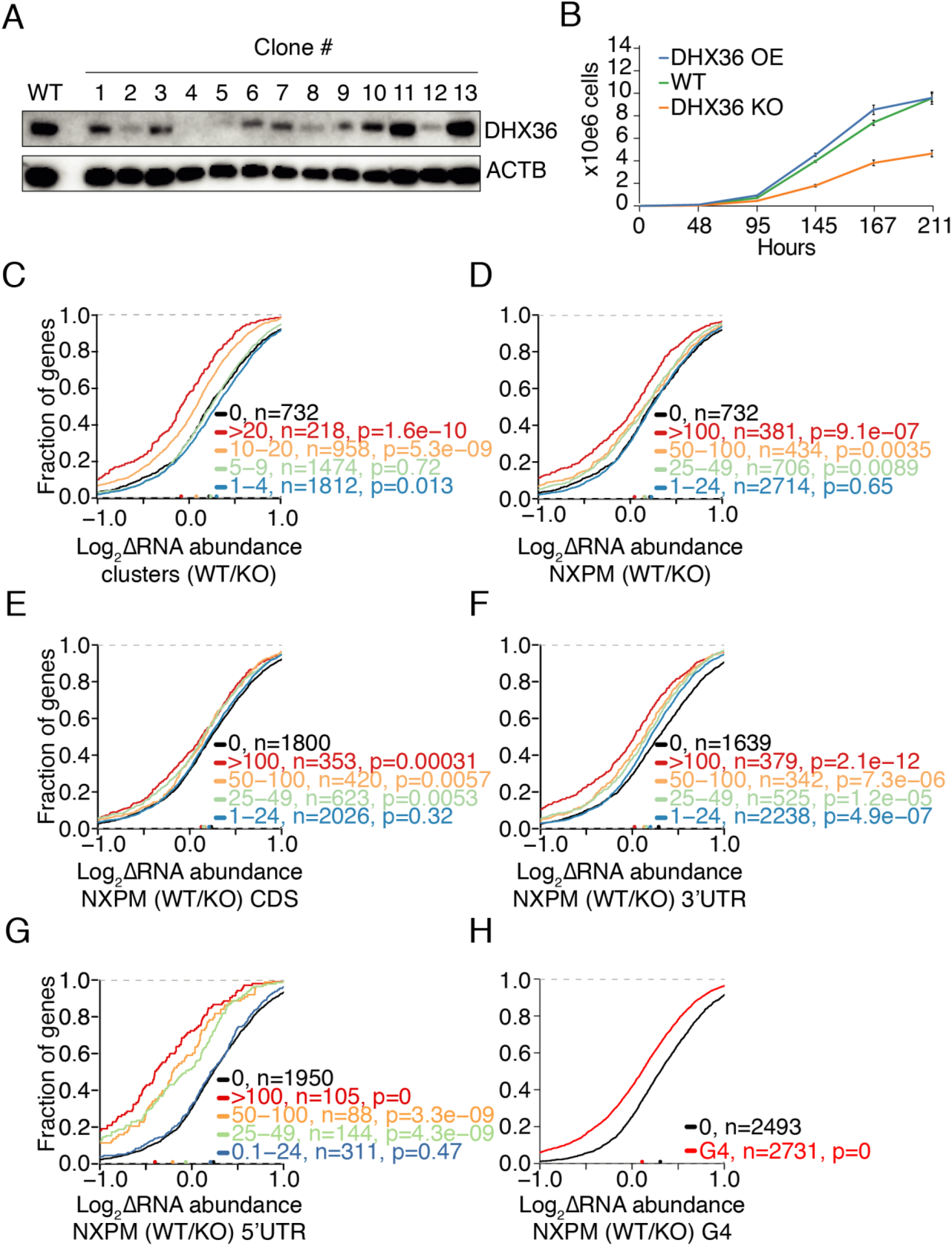
DHX36 KO results in increased target RNA abundance. (**A**) Western blot screening after Cas9-mediated gene editing shows that DHX36 was knocked out in clones 4 and 5. ACTB serves as loading control. (**B**) DHX36 KO (KO) severely reduces cell proliferation compared to wild type (WT). 1×10^5^ cells were seeded at t=0 and the cell number was counted (x10^6^) at indicated timepoints. Error bars represent standard deviations of three independent experiment. (**C**) DHX36 KO results in an increased target mRNA abundance shown by cumulative distribution functions comparing changes in target mRNA abundance of DHX36 knockout cells (n=3) and parental HEK293 cells (n=3). Target mRNAs were binned in accordance to the number of binding clusters obtained by DHX36 E335A PAR-CLIP. Significance was determined using a two-sided Kolmogorov-Smirnov (KS) test. (**D**) Same as in (C), except target mRNAs were binned in accordance to NXPM. (**E**) Same as in (C), except mRNAs were binned based on the number of NXPM in the CDS. (**F**) Same as in (C), except mRNAs were binned based on the number of NXPM in the 3’ UTR. (**G**) Same as in (C), except mRNAs were binned based on the number of NXPM in the 5'UTR. (**H**) Same as in (C), except mRNAs were binned based on whether they harbor a G4-site identified previously (Kwok et al., 2016) overlapping with PAR-CLIP binding sites or not.

Subsequently, DHX36 KO cells where used to investigate the effect of DHX36 on target mRNA abundance using RNA-seq (Suppl. Table S3). For the following analyses we focused on DHX36 targets from the deeper FH-DHX36 E335A PAR-CLIP dataset (Fig. 4). Nevertheless, we observed similar results using the highly overlapping FH-DHX36 PAR-CLIP data (Suppl. Fig. S4). Loss of DHX36 led to an increase in its targets mRNA levels. This effect was dependent on the number of binding sites (Fig. 4C) and to a binding coefficient calculated as the number of crosslinked PAR-CLIP reads per million normalized standardized by target RNA expression (NXPM) (Fig. 4D). We found that both these metrics correlated well with the occupancy of an RBP on its target (Ascano et al., 2012; Hafner et al., 2010). For the top FH-DHX36 E335A targets binned by cluster number (> 20 clusters, n = 218) or NXPM (NXPM > 100, n = 381) mRNA levels were increased upon DHX36 loss by 25% and 15%, respectively.

We further investigated the effect of the DHX36 binding site location on mRNA abundance and binned our targets according to binding in the 3’ UTR, 5´ UTR, or CDS. We found that binding to the UTRs, particularly the 5’ UTR, conferred a considerably stronger effect on mRNA abundance compared to CDS binding sites (Figs. 4E-G). We tried to refine this analysis by identifying mRNAs that were exclusively bound by DHX36 either in the CDS or in the UTRs. Only 130 mRNAs (and only 4 with exclusive binding 5’ UTR) that met these criteria could be identified. Although this analysis was less robust it revealed that CDS binding sites did not influence mRNA abundance. Taken together our data suggest that only UTR sites confer posttranscriptional regulatory effects.

Because of the large overlap of our DHX36 binding sites to potential G4 motifs or other stable structures *in vitro* (Kwok et al., 2016), we asked whether the abundance of mRNAs is affected by DHX36 binding at these special regions. Indeed, target mRNAs bound by DHX36 at potential G4 structures increased in abundance by ~16% in DHX36 KO cells indicating that DHX36 maybe involved in their turnover (Fig. 4H).

### DHX36 increases translational efficiency of its targets

DHX36 was proposed to be involved in translational regulation (Thandapani et al., 2015). We therefore asked whether DHX36 affected translation and directly measured the impact of DHX36 on ribosome occupancy on its targets using ribosome footprinting (Ribo-seq) (Ingolia et al., 2009) in DHX36 KO and the corresponding parental cells (Suppl. Table S4). DHX36 loss resulted in a marginal, albeit statistically significant decrease in ribosome protected fragments (p <10^−5^, two-sided Kolmogorov-Smirnov test, Fig. 5A, B and Suppl. Fig. S5A, B) independent of whether the binding sites were found in the CDS or the UTRs (Fig. 5C-E and Suppl. Fig. S5C, D). We also measured changes in global protein levels using stable isotope labeling in cell culture (SILAC) followed by mass spectrometry. A modest but significant decrease in protein levels from DHX36 top targets upon DHX36 loss was detected, which is consistent with the decrease in RPFs (Fig. 5G and Suppl. Fig. S5F and Suppl. Table S4).

**Figure 5:**
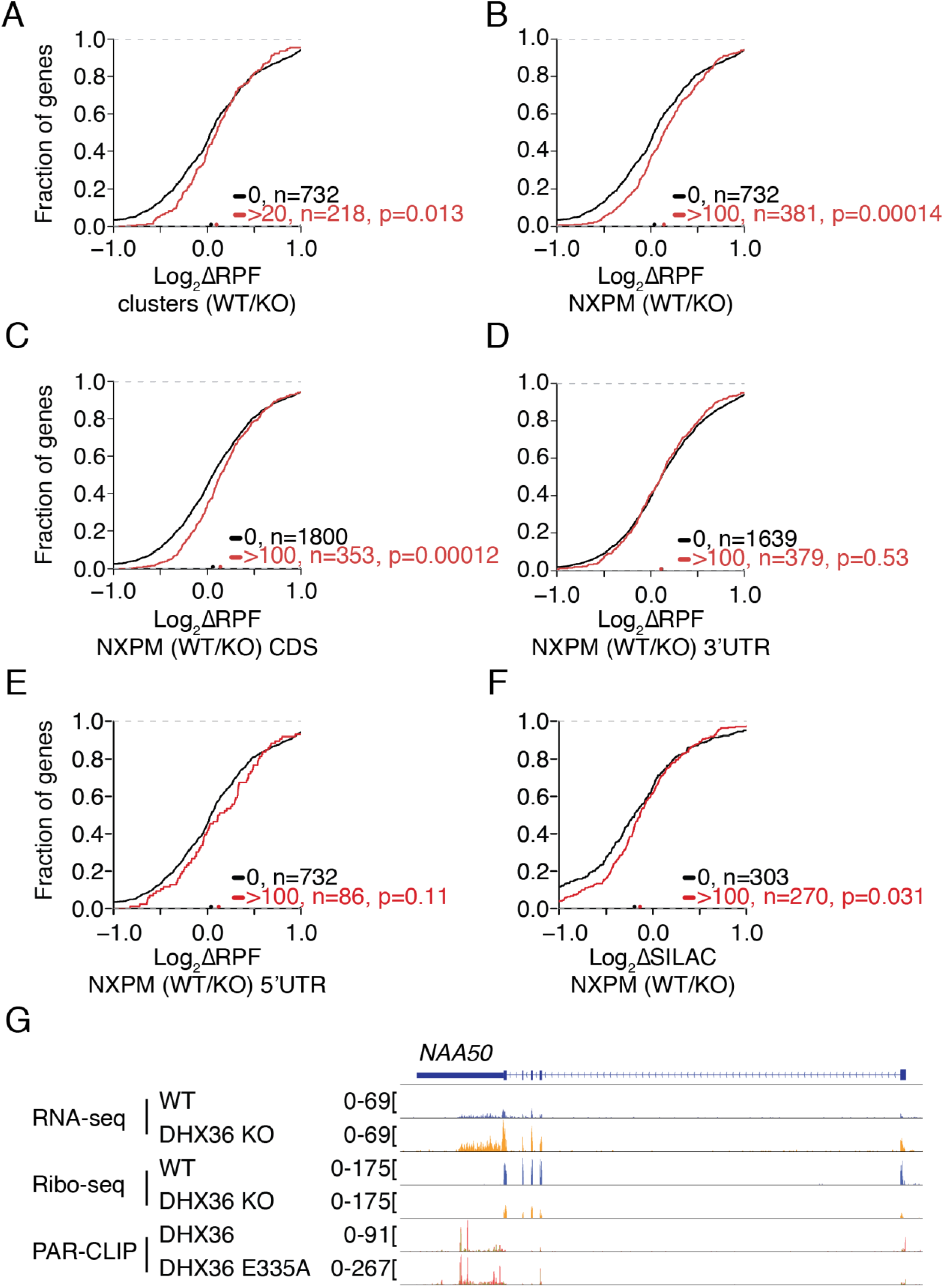
DHX36 KO results in slightly reduced target mRNA translation. (**A**) Cumulative distribution function comparing changes in ribosome-protected fragments (RPFs) of DHX36 KO (n=3) and parental HEK293 cells (n=3). Target mRNAs are binned in accordance to the number of binding clusters obtained by DHX36 E335A PAR-CLIP. Significance was determined using a two-sided KS-test. (**B**) Same as in (A), except target mRNAs were binned in accordance to NXPM. (**C**) Same as in (A), except mRNAs were binned based on the number of NXPM in the CDS. (**D**) Same as in (A), except mRNAs were binned based on the number of NXPM in the 3’ UTR. (**E**) Same as in (A), except mRNAs were binned based on the number of NXPM in the 5’ UTR. (**F**) Same as in (A), except protein abundance changes as determined by SILAC were plotted. **(G)** Screenshot of RNA-seq and Ribo-seq coverage in wildtype and DHX36 KO HEK293 cells on the representative DHX36 target NAA50. Bottom two tracks show the coverage for DHX36 and DHX36 E335A PAR-CLIP.

We calculated the average density of ribosomes on each mRNA in DHX36 KO and control cells by normalizing the number of RPFs with the mRNA abundance. This score, known as the translational efficiency (TE), removes effects of mRNA abundance and approximates the translational output for each mRNA molecule of a given gene (Bazzini et al., 2012; H. Guo et al., 2010; Ingolia et al., 2009). DHX36 KO strongly correlated with a decreased TE on DHX36 targets, independent of binning by PAR-CLIP binding site number or NXPM (~27% decrease for the 385 and 506 top DHX36 targets with NXPM > 100 or > 5 binding sites, respectively (Fig. 6 A, B and Suppl. Fig. S6). Interestingly, the decrease in target TE upon DHX36 KO was again more pronounced in targets bound in the UTRs than in the CDS (Fig. 6C-E and Suppl. Fig. S6). Analogous to RNA abundance shown above, potentially G4 forming RNAs exhibited a 17% decreased TE upon DHX36 KO (Fig. 6F). Taken together with our observation that >90% of DHX36 were not co-sedimenting with translating ribosomes and thus was unlikely to influence translation elongation, our data suggest that DHX36 increases the translational competence of mRNAs by resolving potentially G4 forming structures in 3' and 5’ UTRs and thereby allowing access for the translation initiation machinery.

**Figure 6:**
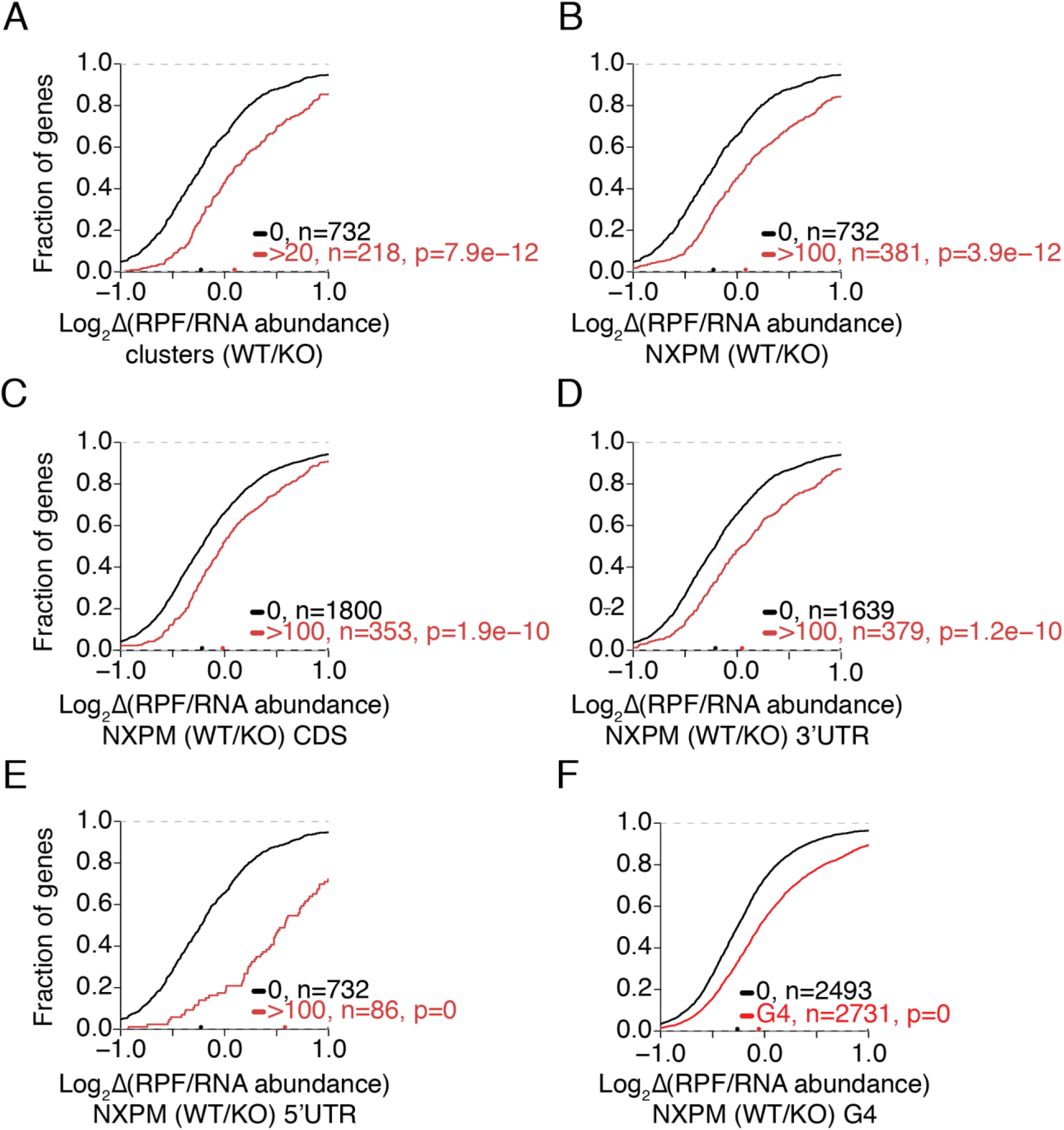
DHX36 increases target mRNA translational efficiency. (**A**) Cumulative distribution function comparing changes in translation efficiency (TE, RPF/RNA abundance) of DHX36 KO (n=3) and parental HEK293 cells (n=3). Target mRNAs are binned in accordance to the number of binding clusters obtained by DHX36 E335A PAR-CLIP. Significance was determined using a two-sided KS test. (**B**) Same as in (A), except target mRNAs were binned in accordance to normalized crosslinked reads per million. (**C**) Same as in (A), except mRNAs were binned based on the number of NXPM in the CDS. (**D**) Same as in (A), except mRNAs were binned based on the number of NXPM in the 3’ UTR. (**E**) Same as in (A), except mRNAs were binned based on the number of NXPM in the 5’ UTR. (**F**) Same as in (A), except mRNAs were binned based on whether they harbor a G4-site identified previously (Kwok et al., 2016) overlapping with PAR-CLIP binding sites or not.

### DHX36 posttranscriptionally regulates mRNA stability

Its mainly cytoplasmic localization suggests a posttranscriptional mechanism of action for DHX36. Nevertheless, isolated reports pointed at a function in DNA G4 recognition and thus transcriptional regulation (Huang et al., 2012). We formally investigated whether the accumulation of target transcripts in DHX36 KO was due to an increase in their transcription or their stability. We isolated nascent chromatin-associated RNA in wildtype and DHX36 KO cells followed by RNA sequencing. Compared to wildtype cells DHX36 KO cells showed no increase in newly-synthesized target mRNAs (Fig. 7A and Suppl. Table S5), suggesting no changes in transcriptional output. We also blocked transcription with actinomycin D and collected samples at different time points after treatment to measure mRNA turnover. All four representative DHX36 targets that we quantified by qPCR had an increased half-life in DHX36 KO cells (Fig. 7B, C and Suppl. Fig. S7A, B). Taken together with our RNA-seq data from DHX36 wildtype and KO cells, these results demonstrate that target mRNA turnover is promoted by DHX36, and that the formation of G-rich structures in their UTRs prevent their degradation. To check if G4 structures themselves have an effect on mRNA stability, we analyzed mRNA abundance by RNA-seq after the addition of 2 μM of the G4 stabilizing small molecule PDS for 42 h. G4 stabilization by PDS in wildtype cells resulted in a significant increase in DHX36 target mRNA abundance, even exceeding the effect of DHX36 loss (Fig. 7D and Suppl. Table S5) and suggesting that formation of G4 structures indeed reduced mRNA turnover or prevented degradation.

**Figure 7:**
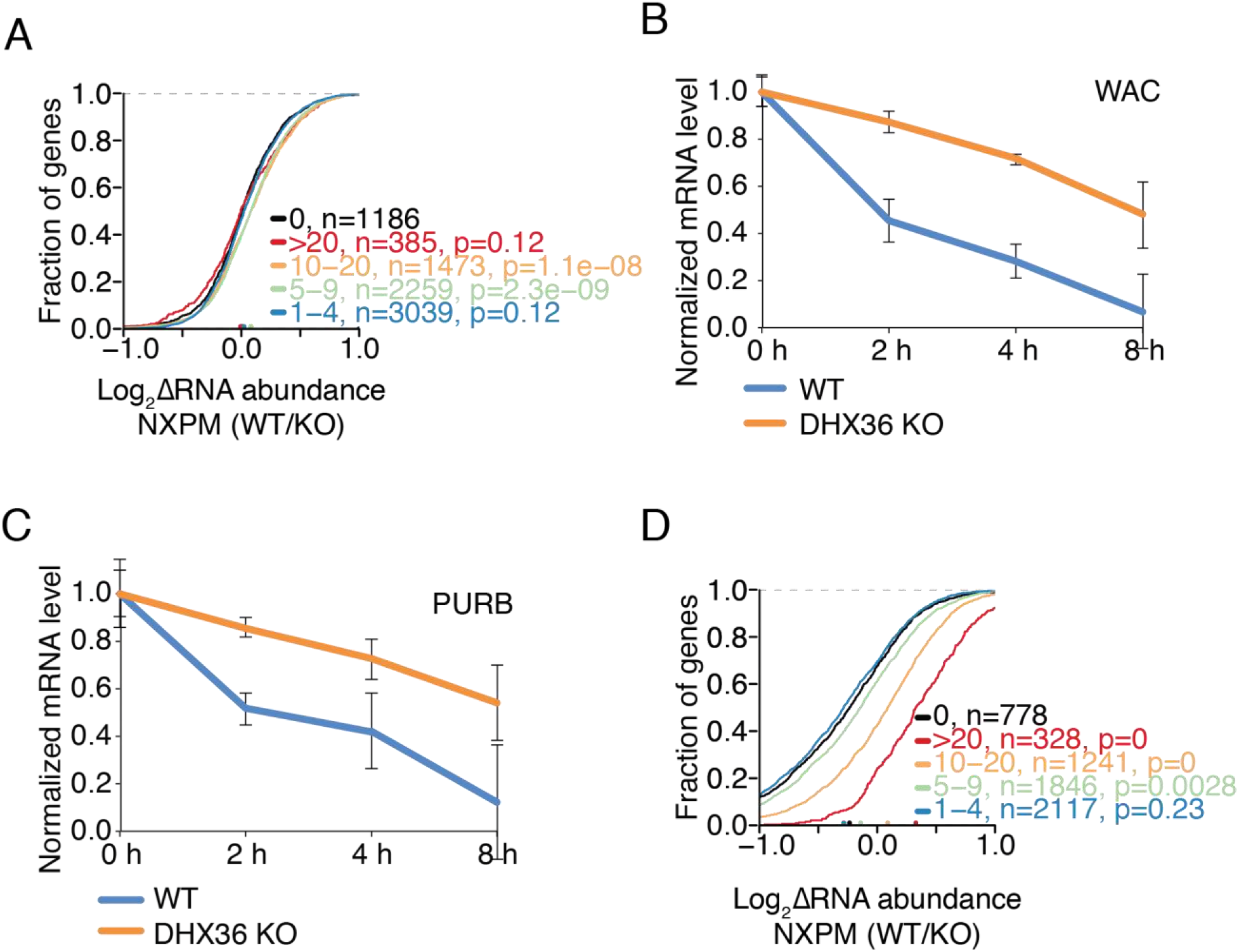
DHX36 regulates target mRNA stability by resolving G-rich structures in their UTRs. **(A)** DHX36 KO regulates target mRNA abundance at a posttranscriptional rather than transcriptional level shown by cumulative distribution functions comparing changes in nascent target mRNA abundance purified from chromatin of DHX36 knockout cells (n=3) and parental HEK293 cells (n=3). Target mRNAs were binned in accordance to the number of binding clusters obtained by DHX36 E335A PAR-CLIP. Significance was determined using a two-sided KS test. **(B,C)** DHX36 target mRNAs increase their half-life upon DHX36 KO shown by qPCR of WAC (B) and PURB (C) after transcriptional block using Actinomycin D and isolation of RNA at the indicated timepoints. **(D)** Stabilization of G4 structures using pyridostatin (PDS) in HEK293 cells results in the accumulation of DHX36 PAR-CLIP targets to a larger degree than DHX36 KO shown by cumulative distribution functions comparing changes in target mRNA abundance of DHX36 knockout cells (n=3) and parental HEK293 cells treated wit PDS (n=3). Target mRNAs were binned in accordance to the number of binding clusters obtained by DHX36 E335A PAR-CLIP. Significance was determined using a two-sided KS test.

## DISCUSSION

Here we present a comprehensive and systems-wide characterization of the targets and function of the DEAH-box helicase DHX36. We identified target RNA binding sites transcriptome-wide, delineated consensus binding motifs, and globally defined the effect of DHX36 binding on target mRNA abundance and translation. To date only few DHX36 targets were described, and our approach expanded this list and demonstrated that DHX36 is specifically interacting with thousands of RNAs, including 22 of the 27 previously known ones (Booy et al., 2014; Nie et al., 2015; Sexton and Collins, 2011; Tran et al., 2004) (Suppl. Table S1). We propose a model in which DHX36 disrupts structured regions in mRNA UTRs, which would otherwise render the mRNA translation incompetent and prevent their turnover.

Two seemingly antagonistic *in vivo* binding motifs were offered for DHX36: on the one hand it bound A/U-rich elements in the 3’ UTR of the PLAU mRNA and recruited destabilizing factors, such as PARN (Tran et al., 2004), and on the other hand it interacted with DNA or RNA G4 structures unwinding them *in vitro* (Creacy et al., 2008; Vaughn et al., 2005). Mapping DHX36 binding sites transcriptome-wide allowed us to reconcile these binding models (Fig. 2). DHX36 did indeed preferentially bind G-rich sequences, including sites in the 3’ UTR of the first described DHX36 target gene, PLAU. Nevertheless, we also observed an enrichment of A/U-rich sequences in DHX36 PAR-CLIPs using the catalytically inactive mutant DHX36 E335A (Fig. 3). We think A/U-rich elements are genuine binding sites of DHX36 and may even serve as the protein recruitment sites to mRNAs. However we suggest that DHX36 translocation from these sites is rapid due to their less structured nature, which prevents efficient UV crosslinking to the catalytically active helicase.

Most hypotheses on DHX36 function relate to its ability to bind and resolve G4 structures. The top binding sites recovered by PAR-CLIP fulfill the rules for G4 formation and PAR-CLIP recovered more than 50% of the G4 structures formed *in vitro* on HeLa mRNA (Kwok et al., 2016) underscoring the connection between DHX36 and G4 structures (Figs. 4, 5). Nevertheless, we identified additional G-rich regions with strong DHX36 binding that are not necessarily predicted to form canonical G4 structures. These sequence stretches may belong to recently reported G4 classes with less guanines and longer loops, which complicates their prediction (Pandey et al., 2013). Alternatively, DHX36 could be involved in the resolution of other structured regions in the transcriptome.

In eukaryotes, an entire machinery likely exists that suppresses G4 and other structures, considering that on one hand G4 structures are globally unfolded and on the other hand G4 forming RNAs are toxic in prokaryotes that do not contain predicted G4 structures in their transcriptome (J. U. Guo and Bartel, 2016). Using DMS-seq, Guo et al. also found that neither depletion of ATP, nor of DHX36 itself resulted in the detection of G4 *in vivo*, which they interpreted as helicase-independent G4 resolution by RBPs. However, currently at least four helicases are known to affect RNA G4 unfolding *in vitro* (Sauer and Paeschke, 2017) and these proteins possibly compensate for each other's loss, dampening the effect of the knockdown of a single factor. For example, on DNA G4 structures, Rrm3 rescues for the loss of Pif1 helicase (Paeschke et al., 2013). The fact that DHX36 KO cells remain viable, altough with a growth defect (Fig. 4) and only exhibit an ~30% accumulation of the best DHX36 target mRNAs (Fig. 4) does hint at a larger, compensatory network of G4 resolving factors *in vivo*. Redundancy of the G4 remodeling machinery is also consistent with a recent analysis that showed that the copy numbers of G4 containing RNAs are insufficient to force G4 folding and that only overexpressed RNA will form G4 *in vivo* (Serikawa et al., 2017).

We speculate that most G4 resolving factors remain to be identified, considering that transcriptome-wide functional data are limited to only two RBPs, CNBP and eIF4A, that both have clear preference for target mRNA CDS or 5’ UTR (Benhalevy et al., 2017; Wolfe et al., 2014). For example, the translation initiation factor elF4A is essential for unwinding secondary RNA structures in the 5’ UTR (Jackson et al., 2010; Sonenberg and Hinnebusch, 2009). Its loss results in an increase in ribosome footprints in the 5’ UTR of eIF4A target mRNAs, which was explained by unresolved G4 structures (Wolfe et al., 2014). CNBP, increases translational efficiency by binding and preventing structure formation at G-rich regions in target mRNA CDS (Benhalevy et al., 2017). In contrast to these two examples DHX36 preferentially affected mRNA abundance by binding to UTRs rather than ribosome occupancy, indicating that the factors compensating for DHX36 loss remain uncharacterized. While DHX36 does recognize G4-forming and other G-rich sites in the mRNA CDS, its activity appears not to be required to disrupt them. We predict that other proteins, e.g. CNBP or the ribosome itself (Benhalevy et al., 2017; Endoh and Sugimoto, 2016), resolve those structures.

Until recently, questions focused on the function of RNA G4 motifs in the 5´ UTR, which were shown to impact translation initiation and decrease protein expression by inhibiting the ribosome, using various reporter systems and *in vivo* approaches (Kumari et al., 2007; Song et al., 2016; Wolfe et al., 2014). G4 motifs in 3’ UTRs are less well studied but have been implicated in multiple processes including splicing, polyadenylation, and translation (Beaudoin and Perreault, 2013). Our data suggest that G4 (and other structures targeted by DHX36) in both UTRs have two effects: (1.) they result in the stabilization of the mRNA, and (2.) this stabilization does not result in an increased translation. Translation elongation or termination are unlikely to be affected by DHX36 considering that only a minority of DHX36 is found on polysomes and that we do not find a pile-up of ribosomes close to the stop codons or in the 3’ UTR of target mRNAs. This indicates that the additional target mRNA in DHX36 KO cells is not translation competent, either due to decreased efficiency of translation initiation, or by sequestration of these RNAs into granules, such as P-bodies or stress granules, which are known to recruit G4 structures and function as storage for untranslated mRNA (Byrd et al., 2016; Chalupníková et al., 2008; Hubstenberger et al., 2017; Ivanov et al., 2014). DHX36 might impact translational output by modifying the levels of translatable target mRNAs. Such a regulatory process might be required in highly proliferative cells (e.g. in cancer or during development) with their increased demand on protein synthesis.

## METHODS

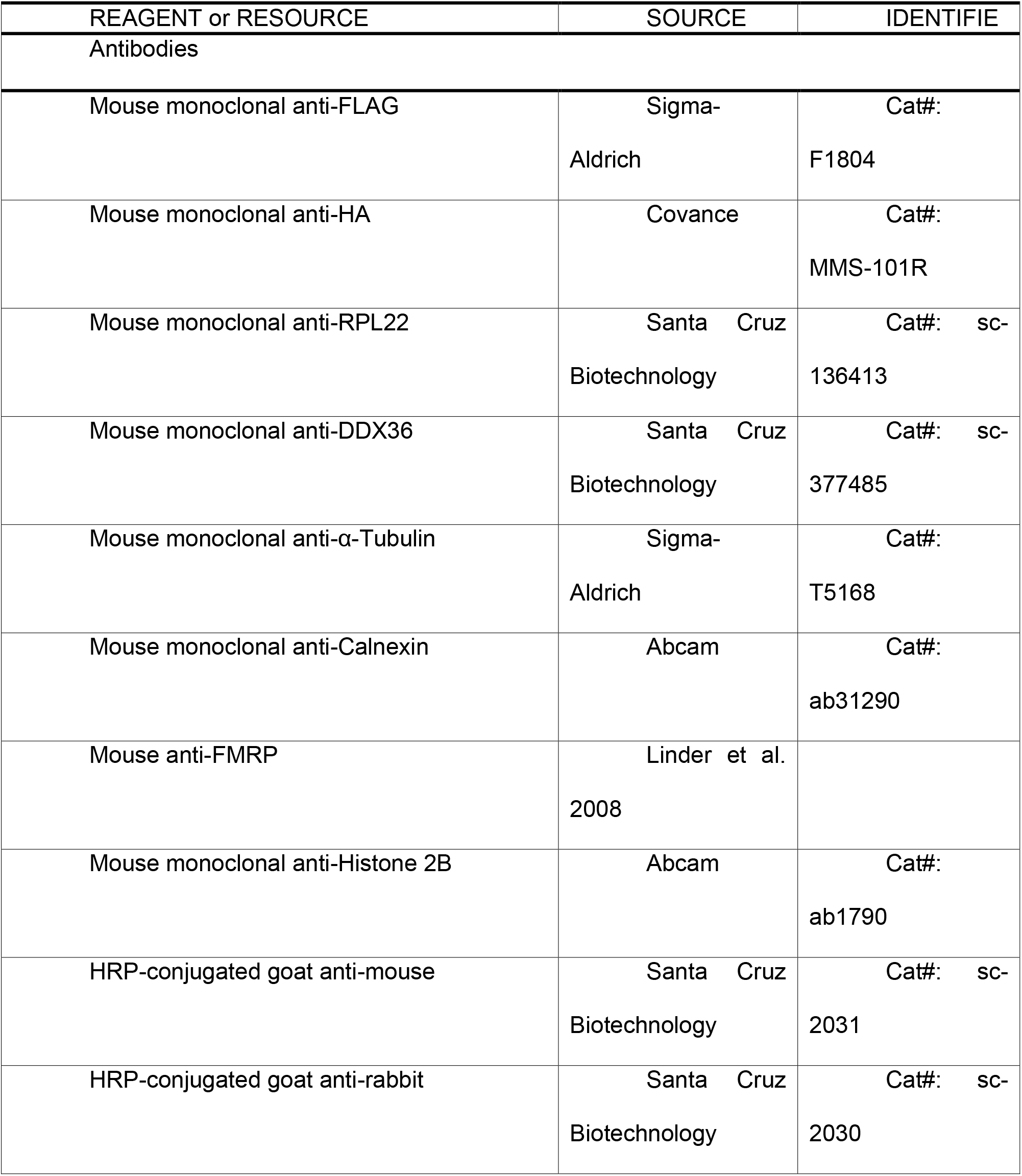
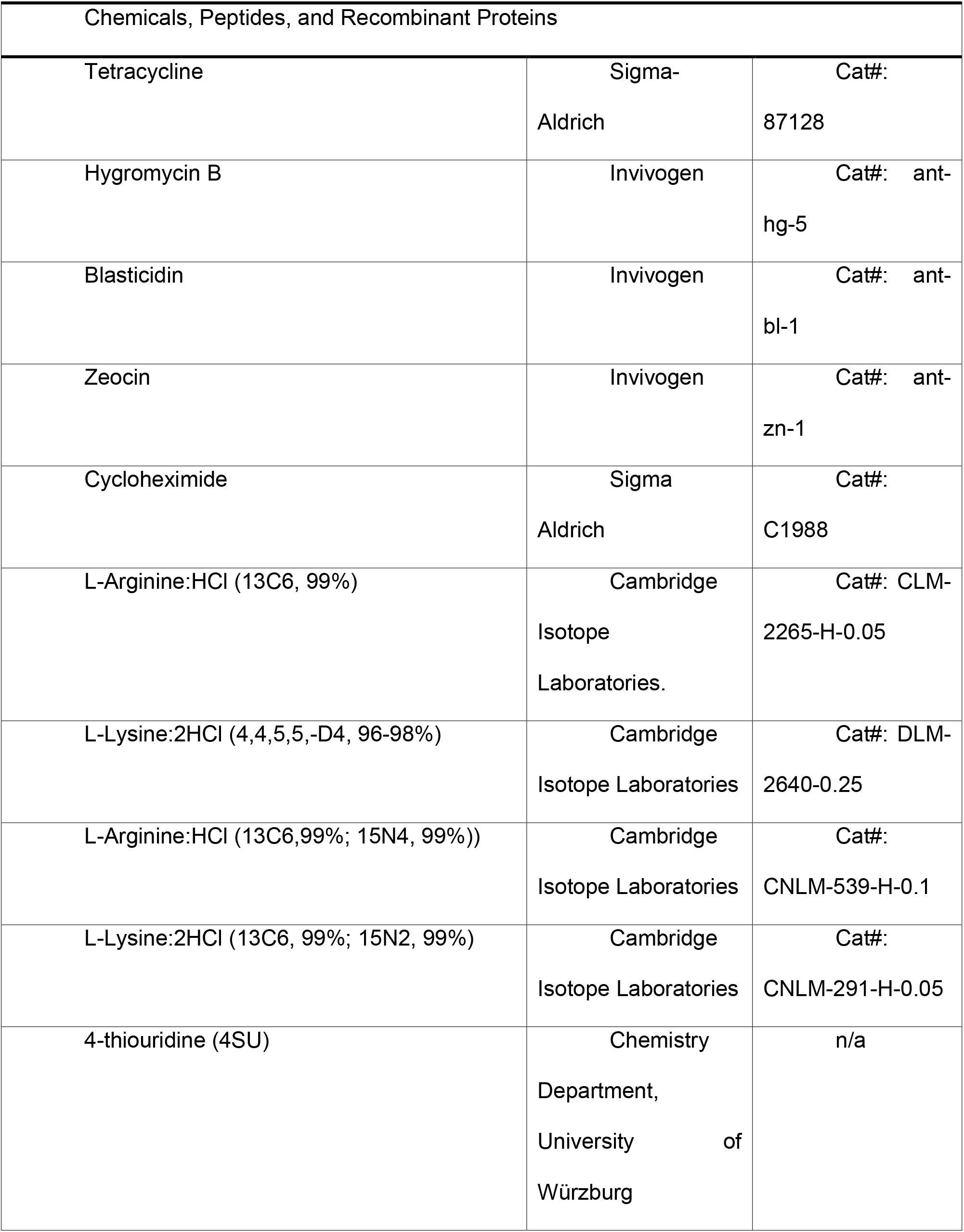
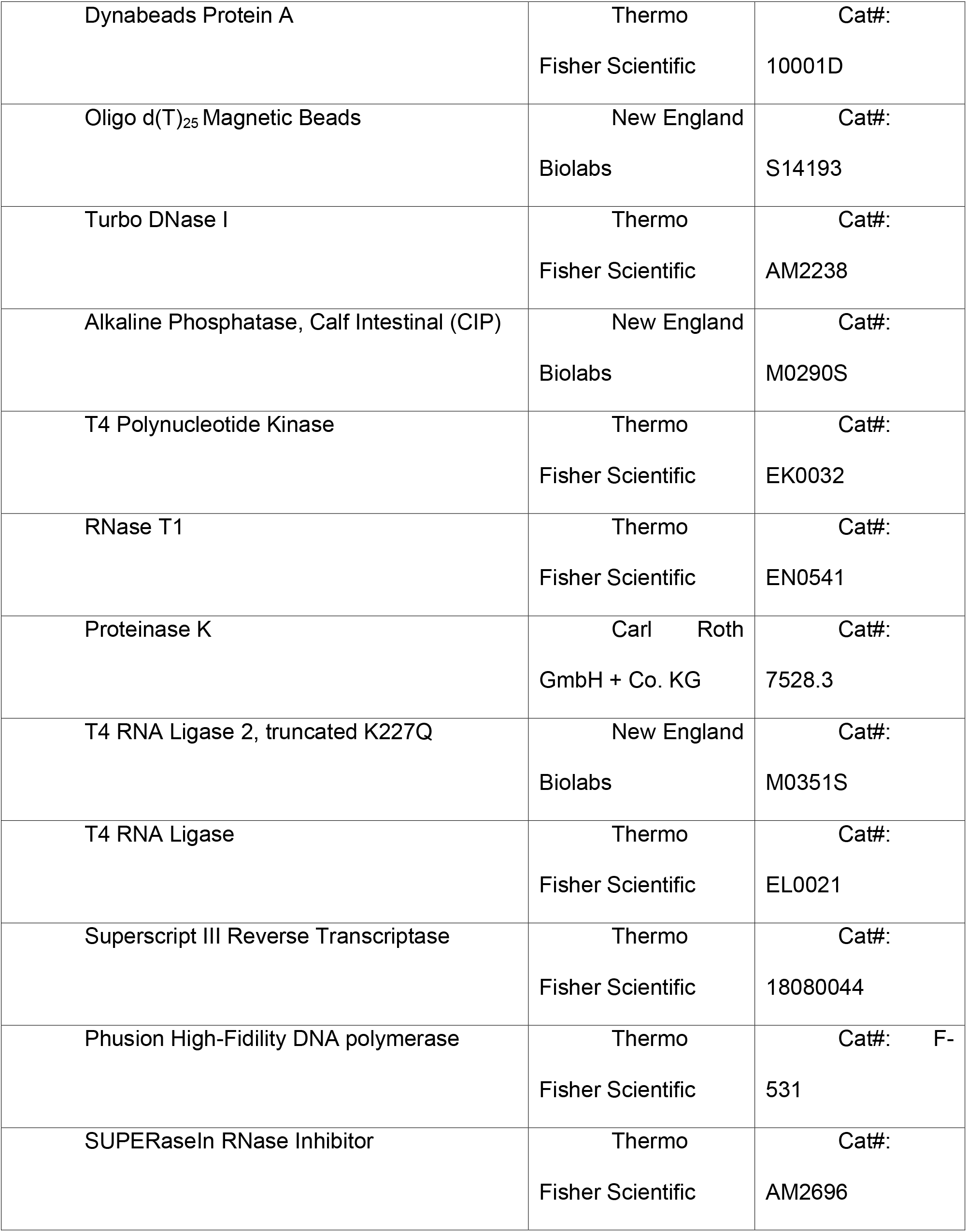
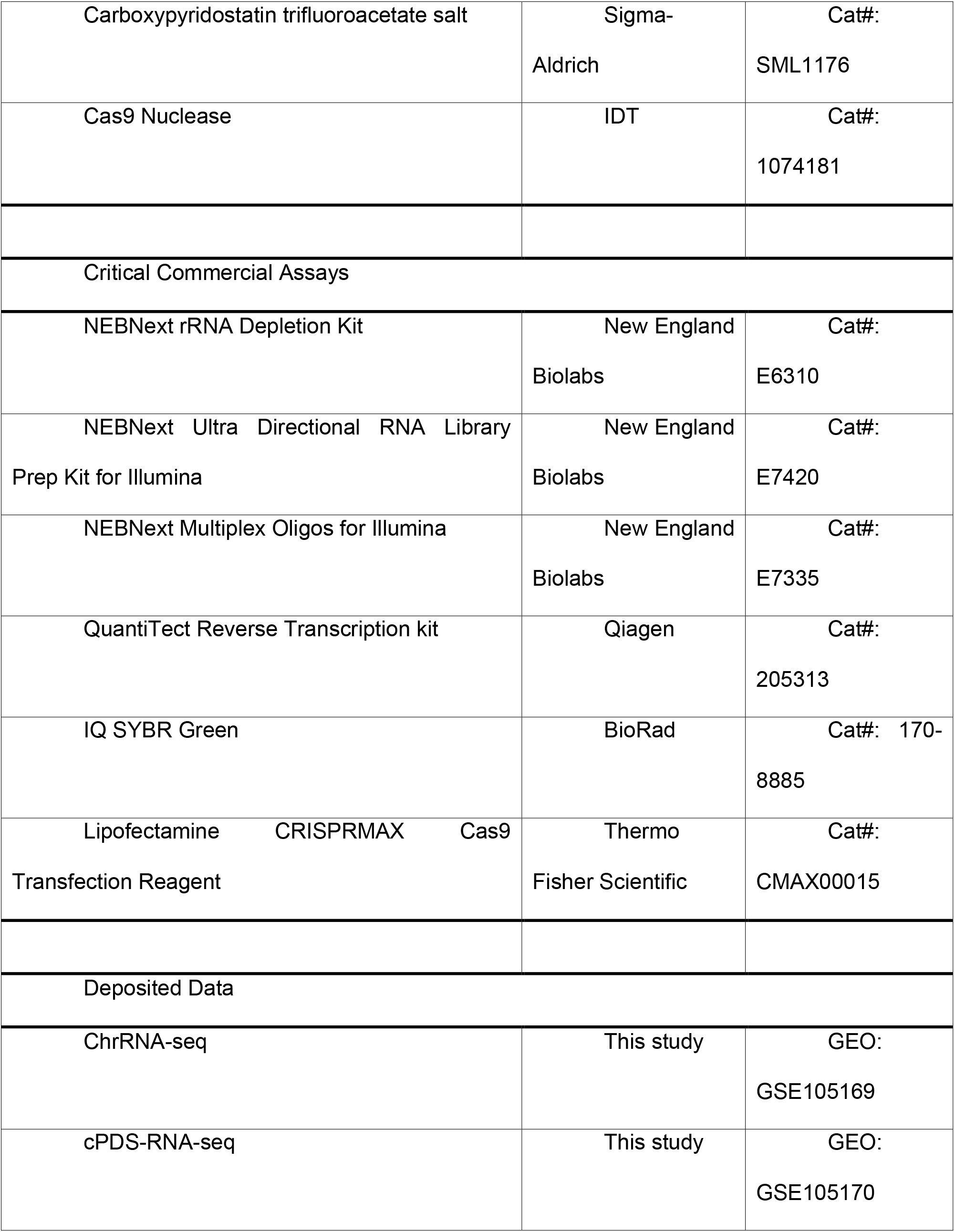
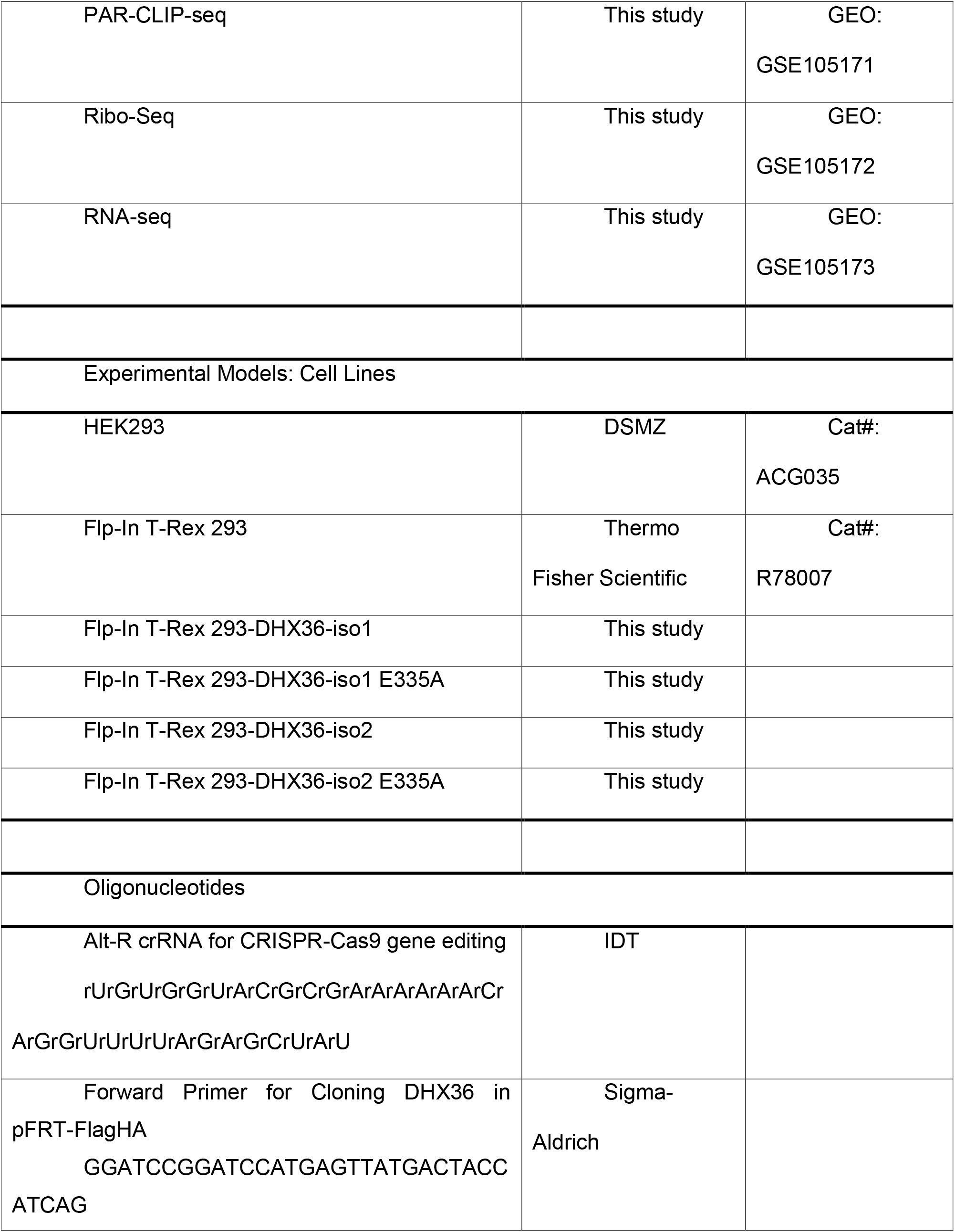
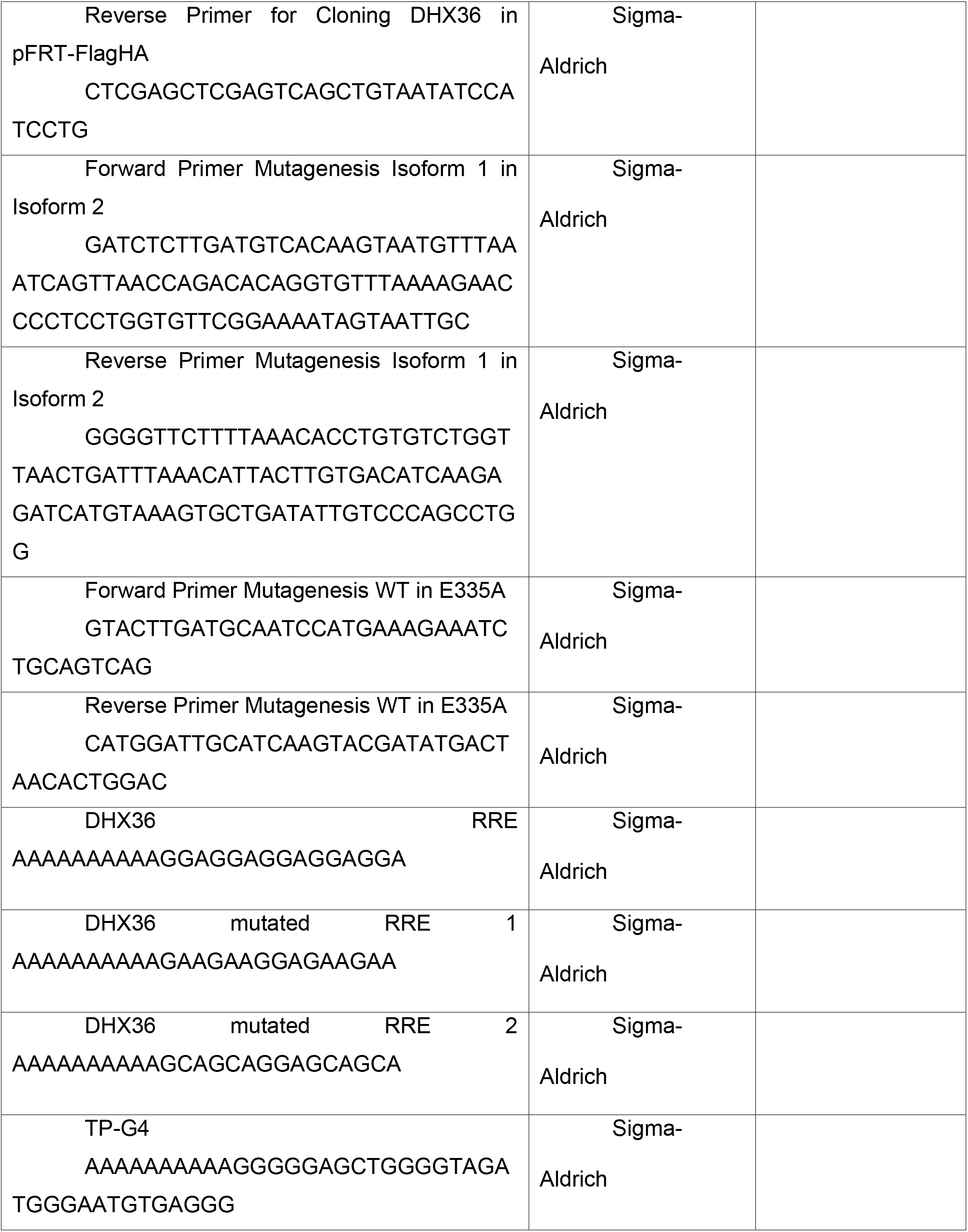
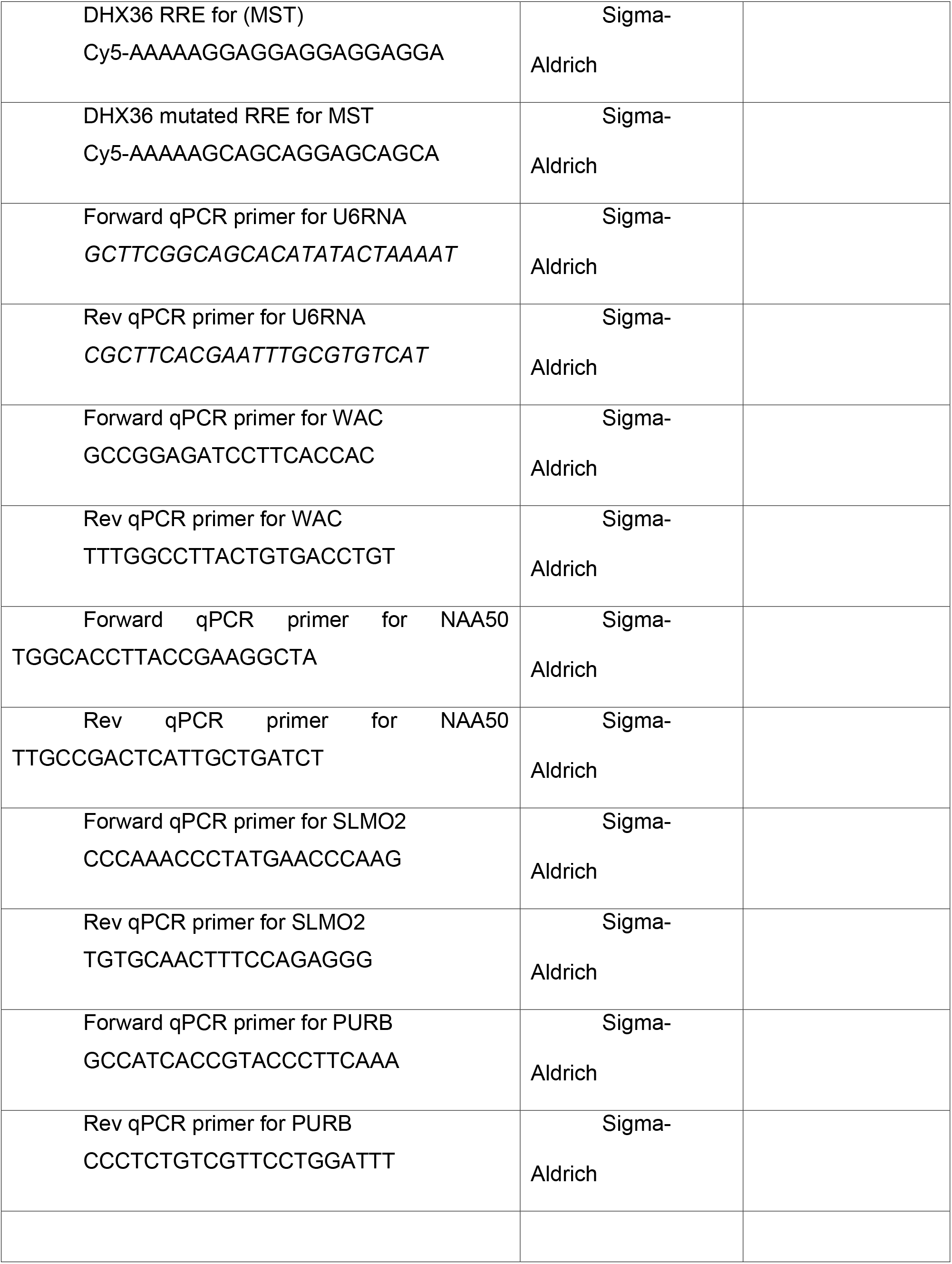
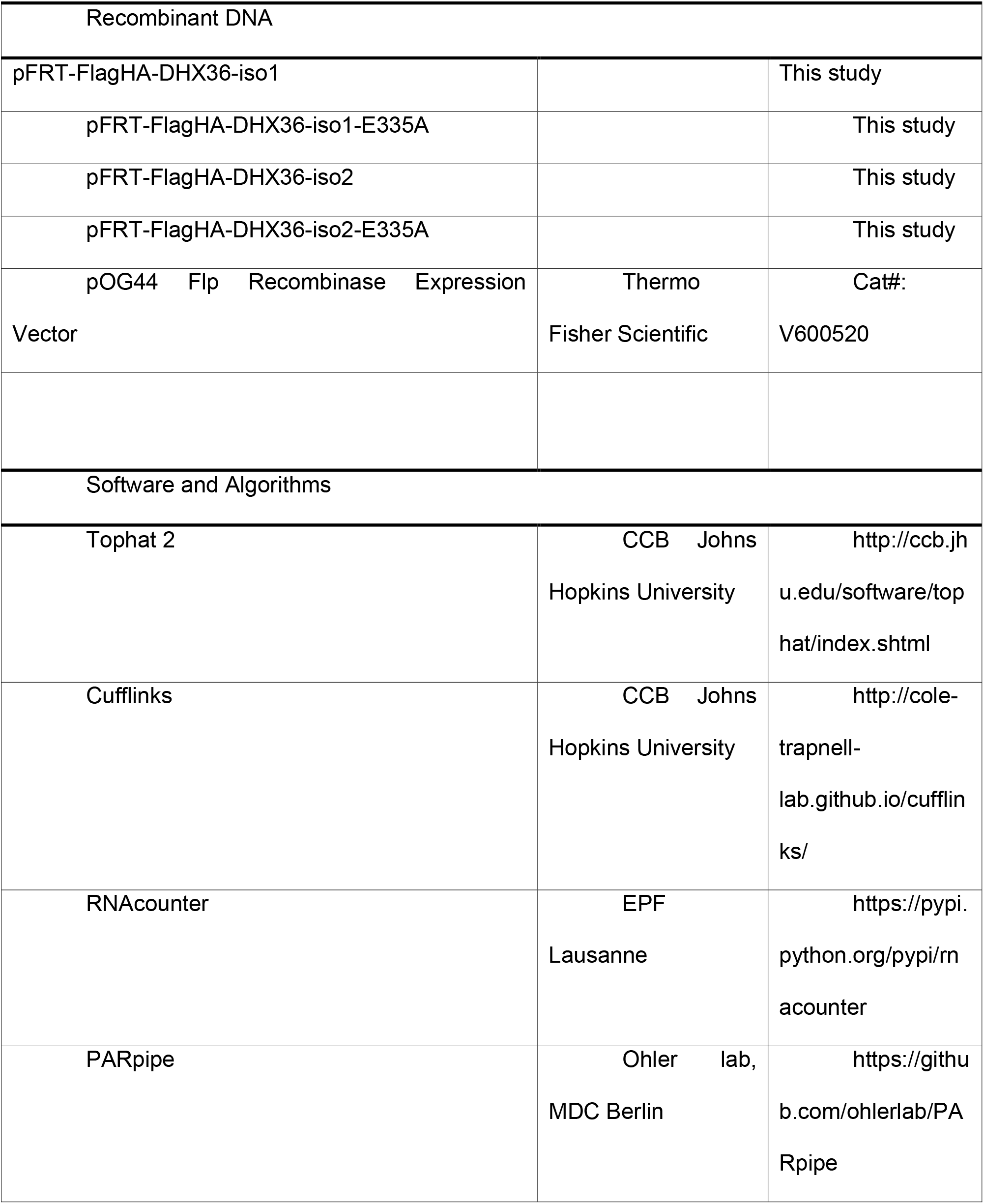
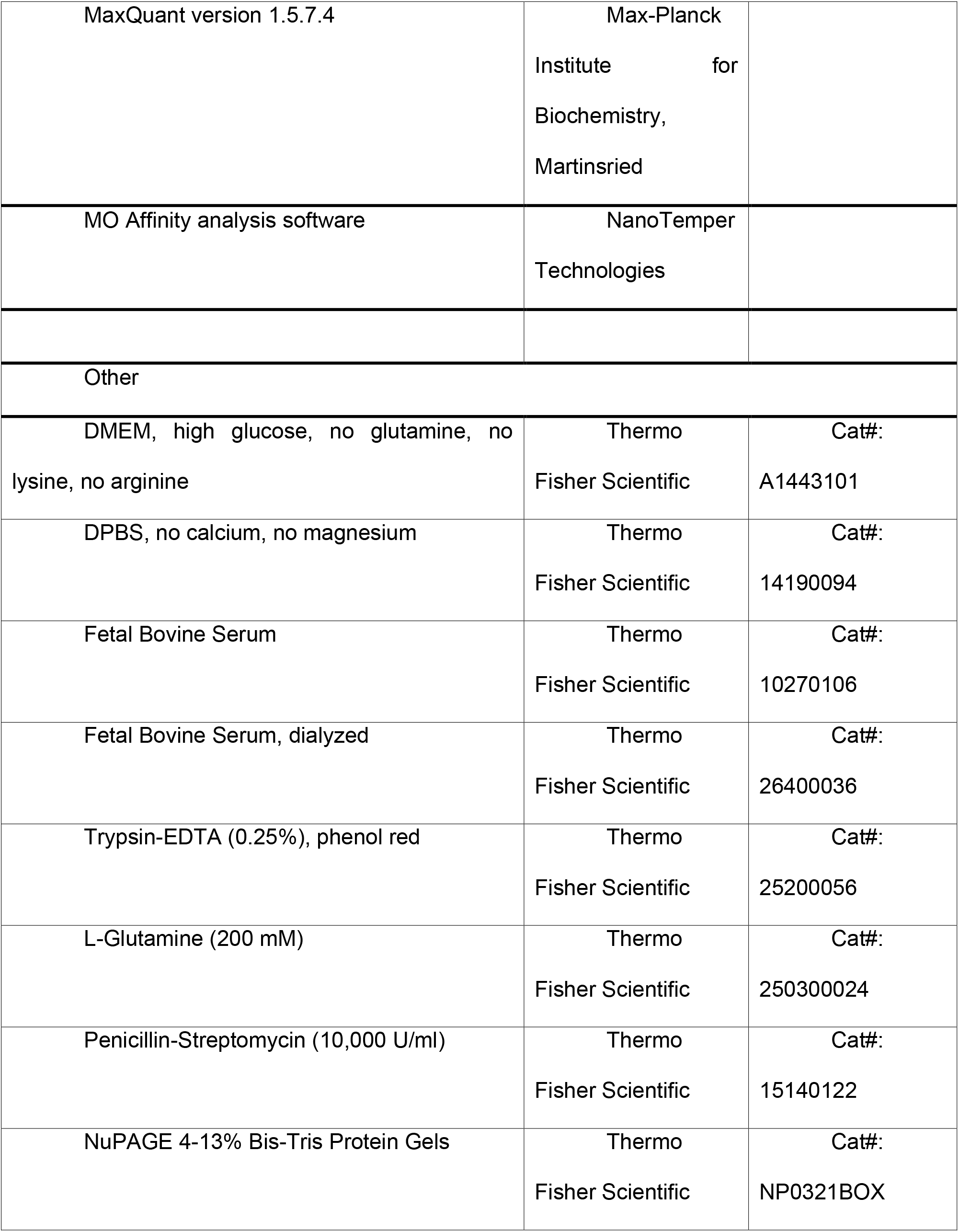

### Cell Lines

HEK293 T-REx Flp-In cells were grown in Dulbecco's modified Eagle's medium (DMEM) supplemented with 10% (v/v) fetal bovine serum (FBS), 100 U ml^−1^ Penicillin-Streptomycin, 100 μg ml^−1^ zeocin, and 10 μg ml^−1^ blasticidin.

### Constructs

Plasmid constructs were generated by standard restriction cloning of a HEK293-cDNA-derived DHX36 insert using BamHI and XhoI. For mutations, standard mutagenesis PCRs were carried out. Used primers are listed in the Resource Table.

### Cell Cultures

Stable Flp-In T-REx HEK293 cell lines inducibly expressing FlagHA-DHX36-Iso1, FlagHA-DHX36-Iso2, FlagHA-DHX36-Iso1-catalytic-dead, or FlagHA-DHX36-Iso2-catalytic-dead were generated as previously described (Spitzer et al., 2013). Essentially, DHX36-Iso1 was cloned from HEK293 cDNA into pFRT/TO/FLAG/HA-Dest plasmid using the restriction enzymes BamHI and XhoI (Landthaler et al., 2008). Based on the so-created pFRT/TO/FLAG/HA-DHX36-Iso1, plasmids for FlagHA-DHX36-Iso2, FlagHA-DHX36-Iso1-catalytic-dead, FlagHA-DHX36-Iso2-catalytic-dead were generated by site-directed mutagenesis. All primer used in this study are listed in the Key Resource Table.

Co-transfection of these plasmid together with pOG44 plasmid using Nanofectin resulted in stable cell lines selected and grown DMEM supplemented with 10% (v/v) FBS, 100 μg ml^−1^ hygromycin B, and 10 μg ml^−1^ blasticidin.

### Subcellular Fractionation

Transgene expression in FlagHa-DHX36-Iso1 and -Iso2 HEK293 cells was induced by addition of 500 ng ml^−1^ tetracycline (Sigma Aldrich) for 15 hours. After washing with ice-cold PBS, induced 100% confluent cells were scrapped off the 14.5-cm cell culture plate using a rubber policeman and collected by centrifugation. Unless otherwise stated, cell fractionation was performed as described by Gagnon et al. (Gagnon et al., 2014). In detail, pelleted cells were resupended in 1 ml of hypotonic lysis buffer (HLB) (10 mM Tris, pH7.5, 10 mM NaCl, 3 mM MgCl2, 0.3% (v/v) NP-40, 10% (v/v) glycerol) per 75 mg cell pellet. After 10 min incubation on ice, cell suspension was briefly vortexed followed by 8 min centrifugation at 800 g and 4°C. The cytoplasmic fraction (supernatant) was thoroughly transferred to a new tube and 5 M NaCl was added to a final concentration of 150 mM. The remaining nuclear fraction (pellet) was carefully washed four times with HLB (and collected after each wash by 2min centrifugation at 200 g and 4°C). After washing, the pellet was resuspended in nuclear lysis buffer (NLB) (20 mM Tris, pH7.5, 150 mM KCl, 3 mM MgCl2, 0.3% (v/v) NP-40, 10% (v/v) glycerol) and sonicated for 2 cycles (40% power, 30 s ON, 2 min OFF). Both the cytosolic and the nuclear fraction were 15 min centrifuged at 18000 g and 4°C to remove all debris. Obtained supernatants were subject of further investigation by standard Western blotting. Used markers for subcellular compartments: nuclear = anti-Histone 2B antibody, cytosolic = anti-α-Tubulin antibody, endoplasmic reticulum membrane = anti-Calnexin antibody.

### RNA-seq

In triplicates, RNA from HEK293 T-REx Flp-In wildtype cells and HEK293 T-Rex Flp-In DHX36 KO cells was isolated with TRIzol reagent according to the manufacturer's instructions. Depletion of ribosomal RNA was accomplished by using the NEBNext rRNA Depletion Kit. cDNA preparation out of resulting RNA was done with the NEBnext Ultra Directional RNA Library Prep Kit for Illumina. cDNA enrichment was facilitated using indexed primers of the NEBNext Multiplex Oligos for Illumina and sequencing was performed on an Illumina HiSeq 2500 platform. Sequencing reads were aligned to the hg19 human genome using Tophat 2 (Trapnell et al., 2012). Cufflinks (Trapnell et al., 2012) was used to quantify reads on the UCSC hg19 annotation set differential expression was determined by Cuffdiff (Trapnell et al., 2012).

### Ribosome footprinting

Ribosome footprinting was performed as published by Ingolia et al (Ingolia et al., 2012). Briefly, triplicates for both HEK293 T-REx Flp-In wildtype cells and HEK293 T-Rex Flp-In DHX36 KO cells were grown to 80% confluency on 10 cm cell culture plates. For inhibition of translation, standard growth media was changed to growth media containing 100 μg/ml cycloheximide (CHX). After 30 sec media was aspirated and plates were immediately cooled down on ice and washed with ice-cold PBS. 400 μl of ribosome footprinting buffer (20 mM Tris, pH7.4, 150 mM NaCl, 4 mM MgCl_2_, 1 mM DTT, 100 μg/ml CHX, 1% NP-40, 25 U/ml Turbo DNase I) were added, cells were scrapped off the plate using a rubber policeman and collected in pre-chilled 1.5 ml microcentrifuge tubes. After 10 min incubation on ice, lysates were triturated by passing 10x through a 26-G-needle and cleared by 10 min centrifugation at 20000 g and 4°C. For digestion of unbound RNA, 300 μl of cell extract was treated with 2.5 U/μl RNase I and incubated at room temperature for 45 min with gentle mixing. RNase digestion was stopped by adding 0.65 U/μl SUPERaseIn and extract was ultracentrifuged through a 900 μl sucrose cushion (20 mM Tris, pH7.4, 150 mM NaCl, 4 mM MgCl_2_, 1 mM DTT, 100 μg/ml CHX, 1 M sucrose, 20 U/ml SUPERaseIn) for 4 hours at 70000 rpm (TLA 100.3 rotor) and 4°C. The supernatant was carefully aspirated and ribosome-containing pellet was resuspendend in 150 μl of ribosome footprinting buffer supplemented with 20 U/ml SUPERaseIn. RNA footprints were purified by phenol-chloroform extraction and precipitated with ethanol. After washing twice with 75% ethanol, the air-dried RNA pellet was resolved in 15 μl DEPC-treated water. Small RNA cDNA libraries for next-generation sequencing were prepared as previously described (Hafner et al., 2012b) with following modifications: firstly, for dephosphorylation 500 ng RNA were incubated 30 min at 37°C with 0.7 U/μl calf intestinal phosphatase (CIP) in 1x CutSmart buffer. Afterwards, samples were separated on a 15% urea-PAGE for exclusion of CIP and selection of 20-35 ribonucleotide long fragments. Extracted, precipitated and washed RNA fragments were ligated with preadenylated, barcoded 3'adapters. Next, RNA was precipitated, washed and phosphorylated at the 5'end with 5 U/μl T4 polynucleotide kinase in 1x T4 DNA ligase buffer (20 μl reaction volume) for 30 min at 37°C. Samples were separated on a 15% urea-PAGE for exclusion of PNK and selection of 3'ligated fragments. Extracted, precipitated and washed RNA was ligated with 5'adapters and samples were separated on a 12% urea-PAGE for selection of 5'and 3'ligated fragments. After reverse transcription and cDNA library enrichment, samples were sequenced on an Illumina HiSeq 2500 platform.

After sequencing the reads were aligned to the human genome version hg19 using TopHat (Trapnell et al., 2012) and quantified on RNA defined in the UCSC hg19 annotation database using Cufflinks (Trapnell et al., 2012). Overlaps of DHX36 cluster and different genomic regions were calculated with BEDTools (Quinlan and Hall, 2010).

## PAR-CLIP

Photoactivatable-Ribonucleoside-Enhanced Crosslinking and Immunoprecipitation (PAR-CLIP) was performed with minor modifications as described previously (Hafner et al., 2010; 2012a). Essential steps are described in the following. For cell lines (DHX36-Iso1/2/Iso1-catalytic-dead-HEK293) cells were grown on 15 14.5-cm cell culture plates to 80% confluency. Induction of transgene expression (addition of 500 ng ml^−1^ tetracycline) was performed for 15 hours as well as feeding the cells with 100 μM of either 4-thiouridin (4SU) or 6-thioguanosin (6SG). After washing with ice-cold PBS cells were crosslinked (irradiation with UV light of 365 nm for 5 min) and scrapped off the plate using a rubber policeman. After pelleting by centrifugation cells were resuspended in 7 ml NP-40 lysis buffer (50 mM HEPES, pH7.5, 150 mM KCl, 2 mM EDTA, 0.5 mM DTT, 0.5% (v/v) NP-40, protease inhibitor cocktail) and incubated on ice for 12 min. Cell lysate was clarified by 15 min centrifugation at 20000 g and 4°C. First RNase T1 digestion (1U/μl) was performed for 15 min at 22°C. 75 μl/ml FLAG-M2 antibody conjugated to magnetic Dynabeads^TM^Protein G were added. Antigen capture was performed for 105 min at 4°C on a rotating wheel. Beads were collected on a magnetic rack and washed 3x with NP-40 lysis buffer. For trimming of the co-captured RNA, a second RNase T1 digestion (10U/μl) was performed for 15 min at 22°C with occasional shaking. 3'ends of the RNA fragments were dephosphorylated using 0.5 U/μl calf intestinal phosphatase for 10 min at 37°C, shaking. RNA was radioactively 5'end-labeled using 1 U/μl T4 polynucleotide kinase and 0.5 μCi/μl ^32^P-γ-ATP for 30 min at 37°C. Almost complete 5'end phosphorylation of RNA was accomplished by adding ATP to a concentration of 100 μM for 5 min. So-treated RNA-protein complexes were separated by SDS-Polyacrylamidgelelectrophorsis (SDS-PAGE, 4-12% NuPAGE). For detection, a blanked storage phosphor screen (GE Healthcare) was exposed to the gel and RNA protein complexes of expected size were excised. After gel elution, protein components of the complexes were digested with 5 mg/ml proteinase K for 2 hours at 55°C. Resulting RNA fragments were isolated by phenol-chloroform-isoamyl alcohol (25:24:1) extraction and subject of a small RNA cDNA library preparation protocol as previously described (Hafner et al., 2012b). Here, a preadenylated, barcoded 3'adapter oligonucleotide (rApp-TAATATCGTATGCCGTCTTCTGCTTG) was used.

So-obtained PAR-CLIP cDNA libraries were sequenced on an Illumina HiSeq 2500 platform. Clusters of overlapping sequence reads mapped against the human genome version hg19 were generated using the PARalyzer software (Corcoran et al., 2011) incorporated into a pipeline (PARpipe; https://ohlerlab.mdc-berlin.de/software/PARpipe_119/) with default settings. Binding sites were categorized using the Gencode GRCh37.p13 GTF annotation (gencode.v19.chr_patch_hapl_scaff.annotation.gtf, http://www.gencodegenes.org/releases/19.html).

### Circular Dichroism

For G-quadruplex formation, 100 μM oligodeoxynucleotides in folding buffer (10 mM Tris, pH7.5, 100 mM KCl, 1 mM EDTA) were incubated for 10 min at 95°C, followed by subsequent passive cooling to RT. CD measurement was performed using a Jasco J-810 spectropolarimeter in 0.2 ml quartz cuvettes. Measurements in the range of 200-350 nm were recorded. Measurements were averaged between ten accumulations with an instrument scanning speed of 10 nm/sec. Oligodeoxynucleotides used in this experiment were purchased from Sigma-Aldrich.

### Growth curves

Regular grown HEK293 T-REx Flp-In wildtype cells and HEK293 T-Rex Flp-In DHX36 KO cells were trypsinized, counted and 0.1 × 10^6 cells were seeded in 35 mm cell culture plates. At indicated timepoints, cells were counted in triplicates using a Fuchs-Rosenthal-hemocytometer.

## SILAC

In duplicates, HEK293 T-REx Flp-In wildtype cells and HEK293 T-Rex Flp-In DHX36 KO cells were grown to approximately 60% confluency on 35 mm cell culture plates. Then, cells were fed with light media (DMEM, 10% dialyzed FBS, 4 mM glutamine, 1.74 mM L-proline, 0.8 mM L-lysine, 0.4 mM L-arginine). 24 hours later, cells were washed twice with pre-warmed PBS and wildtype cells were changed to medium-heavy media (DMEM, 10% dialyzed FBS, 4 mM glutamine, 1.74 mM L-proline, 0.8 mM L-lysine [4,4,5,5-D4], 0.4 mM L-arginine [U-13C6]) and KO cells to heavy media (DMEM, 10% dialyzed FBS, 4 mM glutamine, 1.74 mM L-proline, 0.8 mM L-lysine [U-13C6, 15N2], 0.4 mM L-arginine [U-13C6, 15N4]). 24 hours later, cells were washed twice with pre-warmed PBS and cells were collected in 100 μl NP-40 lysis buffer (50 mM HEPES, pH7.5, 150 mM KCl, 2 mM EDTA, 0.5 mM DTT, 0.5% (v/v) NP-40, protease inhibitor cocktail). After incubation for 10 min on ice, lysates were cleared by 15 min centrifugation at 20000g and 4°C. Protein concentration was assessed by standard Bradford assay.

For in-gel digestion proteins were reduced and alkylated prior to SDS-PAGE by heating the cell lysates for 10 min at 70°C in NuPAGE LDS sample buffer (Life Technologies) supplemented with 50 mM DTT. Samples were alkylated by adding 120 mM iodoacetamide Simply Blue (Life Technologies). Whole lanes were cut into 15 bands. The bands were destained with 30% acetonitrile, shrunk with 100% acetonitrile, and dried in a vacuum concentrator. Digests with 0.1 μg trypsin (Promega) per gel band were performed overnight at 37°C in 50 mM ammonium bicarbonate (ABC) buffer. Peptides were extracted from the gel slices with 5% formic acid. NanoLC-MS/MS analyses were performed on an Orbitrap Fusion (Thermo Scientific) equipped with an EASY-Spray Ion Source or a PicoView Ion Source (New Objective) and coupled to an EASY-nLC 1000 (Thermo Scientific). Using the Easy-Spray Ion Soruce the peptides were loaded on a trapping column (2 cm × 75 μm ID, PepMap C18, 3 μm particles, 100 Å pore size) and separated on an EASY-Spray column (25 cm × 75 μm ID, PepMap C18, 2 μm particles, 100 Å pore size). Using the PicoView Ion Source the peptides were loaded on capillary columns (PicoFrit, 30 cm x 150 μm ID, New Objective) self-packed with ReproSil-Pur 120 C18-AQ, 1.9 μm (Dr. Maisch). A 60-minute linear gradient from 3% to 40% acetonitrile and 0.1% formic acid was used. Both MS and MS/MS scans were acquired in the Orbitrap analyzer with a resolution of 60,000 for MS scans and 15,000 for MS/MS scans. HCD fragmentation with 35% normalized collision energy was applied A Top Speed data-dependent MS/MS method with fixed cycle time of 3 seconds was used. Dynamic exclusion was applied with a repeat count of 1 and exclusion duration of 120 s; singly charged precursors were excluded from selection. Minimum signal threshold for precursor selection was set to 50,000. Predictive AGC was used with an AGC target value of 5e5 for MS scans and 5e4 for MS/MS scans. EASY-IC was used for internal calibration.

For MS raw data file processing, database searches and quantification, MaxQuant version 1.5.7.4 (Cox and Mann, 2008) was used. UniProt human reference proteome database was used in combination with a database containing common contaminants as a reverse concatenated target-decoy database. Protein identification was under control of the false-discovery rate (<1% FDR on protein and peptide level). In addition to MaxQuant default settings, the search was performed with tryptic cleavage specificity with three allowed missed cleaves. The search was performed with the following variable modifications: Gln to pyro-Glu formation and oxidation (on Met). Normalized H/M ratios were used for protein quantitation (at least two peptides per protein

### Polysome profiling

HEK293 T-REx Flp-In wildtype cells were grown on a 14.5 cm cell culture plate to 90-100% confluency. Then, growth media was change to media containing 25 μg/ml cycloheximide. After 10 min incubation, cells were washed once with ice-cold PBS and 100 μl of polysome lysis buffer (20 mM Tris, pH7.5, 100 mM KCl, 5 mM MgCl_2_, 1 mM DTT, 0.5% (v/v) NP-40, 100 μg/ml cycloheximide, 20 U/ml SUPERaseIn, protease inhibitor cocktail) were added (note: for samples used for RNase-treated lysates, no SUPERaseIn was added). Cells were scrapped of the plate using a rubber policeman and transferred to a pre-chilled 1.5 microcentrifuge tube. After 10 min incubation on ice for lysis, lysate was cleared by 10 min centrifugation at 20000 g and 4°C. Clarified lysate was loaded onto a 5-45% linear sucrose gradient (sucrose in 20 mM Tris, pH7.5, 100 mM KCl, 5 mM MgCl_2_) and ultracentrifuged for 60 min in a SW60Ti rotor (Beckman) at 38000 rpm and 4°C. During fractionation performed using a Gradient fractionator (Biocomp) the UV profile (254 nm) was measured. Obtained fractions were further analyzed by standard Western blot.

### Light microscopy imaging

Regular grown HEK293 T-REx Flp-In wildtype cells and HEK293 T-Rex Flp-In DHX36 KO cells were trypsinized, counted and 1.2 × 10^6 cells were seeded in 60 mm cell culture plates. Microscopic images were taken 24, 48, 72, and 96 hours after seeding using a EVOS FL cell imaging system.

### CRISPR/Cas9 Gene Editing

crRNAs were designed using <u>https://benchling.com</u>. Alt-R crRNA was ordered from IDT. 100 pmol Alt-R crRNA and 100 pmol Alt-R tracrRNA-ATTO 550 were denatured at 95˚C for 5 min and incubated at RT for 15 min to anneal both strands in a total volume of 100 μl in Nuclease-Free Duplex Buffer (IDT). 15 pmol annealed RNA was combined with 16 pmol Cas9 (IDT) and 5 μl Cas9+ reagent (Invitrogen) in Opti-MEM (Gibco) in a total volume of 150 μl and mixed well. In a second tube 125 μl Opti-MEM was combined with 7.5 μl CRISPRMAX (Invitrogen) and mixed well. After incubation at RT for 5 min the tubes were combined, mixed well, and formed complex was transferred to a 6-well compartment containing HEK293T cells. The HEK293T cells were seeded the previous day at a density of 4 × 10^5^ cells/ml. After 48 hr ATTO 550 positive cells were FACS sorted and seeded at the density of up to 1 cell per well in a 96 well plate using a standard medium. Single clones were expanded and analyzed for loss of protein by Western blot using anti-DHX36 antibodies.

### cPDS-seq

HEK293 T-REx Flp-In wildtype cells were grown to approximately 50% confluency on 35 mm cell culture plates. Then in triplicates, growth media was changed to standard media or media containing 2 μM carboxy-pyridostatin. After 42 h incubation, cells were collected and total RNA was isolated using TRIzol according to the manufacturer's instructions. Depletion of ribosomal RNA was accomplished by using the NEBNext rRNA Depletion Kit (NEB). cDNA preparation out of resulting RNA was done with the NEBnext Ultra Directional RNA Library Prep Kit for Illumina (NEB). cDNA enrichment was facilitated using indexed primers of the NEBNext Mulitplex Oligos for Illumina (NEB) and sequencing was performed on an Illumina HiSeq 3000 platform. Sequencing reads were aligned to the hg19 human genome using Tophat 2. quantified on RNA defined in the UCSC hg19 annotation database using Cufflinks (Trapnell et al., 2012).

### Isolation of chromatin-associated RNA

In triplicates, standardized cultured HEK293 T-REx Flp-In wildtype cells and HEK293 T-Rex Flp-In DHX36 KO cells were grown on 14.5 cm plates to 80-90% confluency. After collecting the cells by centrifugation. Obtained pellets were washed once with ice-cold PBS and transferred to pre-chilled 1.5 ml microcentrifuge tube. Cells were resuspended in 400 μl ice-cold Cytoplasmic Fraction Buffer (20 mM HEPES, pH7.6, 2 mM MgCl_2_, 10% (v/v) glycerol, 0.5 mM DTT, 0.1% (v/v) NP-40, protease inhibitor cocktail, 40 U/ml Murine RNase Inhibitor (NEB)) and incubated on ice for 5 min. Crude lysates were layered on a 400 μl sucrose buffer cushion (10 mM HEPES, pH7.6, 10 mM NaCl, 1.5 mM MgCl_2_, 10% (v/v) glycerol, 37% (w/v) sucrose, 0.5 mM EDTA; 0.5 mM DTT, protease inhibitor cocktail, 40 U/ml Murine RNase Inhibitor (NEB)) and centrifuged for 20 min at 20000 g and 4°C. Resulting supernatant, representing the cytoplasmic fraction was collected and the pellet was resuspendend in 200 μl ice-cold Nuclear Lysis Buffer (10 mM HEPES, pH7.6, 100 mM NaCl, 50% (v/v) glycerol, 0.5 mM EDTA; 0.5 mM DTT, protease inhibitor cocktail, 40 U/ml Murine RNase Inhibitor (NEB)). 200 μl ice-cold 2x NUN-Buffer (50 mM HEPES, pH7.6, 600 mM NaCl, 7.5 mM MgCl_2_, 2 M urea, 0.2 mM EDTA; 0.5 mM DTT, 2% (v/v) NP-40) were added dropwise, followed by a pulsed vortexing and incubation on ice for 20 min. After 30 min centrifugation at 20000 g and 4°C, resulting supernatant (nucleoplasmic fraction) was collected and the pellet (chromatin), after washing with 1 ml Nuclear Lysis Buffer, was resuspended in 1 ml Trizol. Solutions were heated to 65°C and run through a G26-needle until no insoluble objects were visible. Chromatin-associated RNA was isolated according to the manufacturer's instructions for Trizol-based RNA isolation. cDNA preparation out of resulting RNA was done with the NEBnext Ultra Directional RNA Library Prep Kit for Illumina (NEB). cDNA enrichment was facilitated using indexed primers of the NEBNext Mulitplex Oligos for Illumina (NEB) and sequencing was performed on an Illumina HiSeq 3000 platform. Sequencing reads were aligned to the hg19 human genome using Tophat 2. quantified on RNA defined in the UCSC hg19 annotation database using Cufflinks (Trapnell et al., 2012).

### Actinomycin D mRNA stability assay

HEK293 T-REx Flp-In wildtype cells and HEK293 T-Rex Flp-In DHX36 KO cells were grown on 35 mm plates to 80% confluency. At timepoint T=0, media was changed to media containing 3 μg/ml Actinomycin D. Cells were collected by trypsinization on timepoints 0 h, 2 h, 4 h, and 8 h after addition of Actinomycin D. Total RNA was isolated using TRIZOL according to the manufacturer's instructions. Isolated RNA was reverse-transcribed using the QuantiTect Reverse Transcription Kit (Qiagen). Quantification of target mRNA levels was done by qPCR using gene specific primer on a CFX96 Real Time System with a C1000 Touch Thermal Cycler (Biorad). For normalization primer specific for U6 RNA was used. Data were analyzed using the CFX Manager Software (Biorad).

### Purification of FH-DHX36

Expression of the FH-DHX36 transgene was induced 36 h before harvesting with 500 ng ml^−1^ tetracycline. After washing 3x with ice-cold PBS, cells were and scrapped off the plate using a rubber policeman and collected by centrifugation, followed by 10 min lysis on ice with NP40 lysis buffer (50 mM HEPES, pH 7.5, 150 mM KCl, 2 mM EDTA, 0.5 mM DTT, 0.5% (v/v) NP-40, protease inhibitor cocktail). Obtained crude lysate was run 10x through a G26-needle and insoluble cell debris were removed by centrifugation. In lysis buffer calibrated Anti-Flag M2 Magnetic beads (Sigma) were added and incubated for 5 h on a head-over-tail wheel at 4°C. Beads were collected on a magnetic rack and washed 3x with IP wash buffer (50 mM Tris-HCl, pH 7.5, 300 mM NaCl, 0.1% NP40, 5 mM EDTA). FH-DHX36 was eluted with 0.1 mg/ml FLAG peptide in elution buffer (10 mM Tris-HCl, pH 7.5 and 150 mM NaCl).

### Microscale thermophoresis (MST)

For microscale thermophoresis, binding reactions were prepared in 1x MST buffer, supplemented with 0,5% BSA and 5 mM DTT in a total volume of 40 μl. A constant concentration of 25 nM 5'Cy5 labeled oligonucleotides (folded) were used (see Resource Table). A 1:1 serial dilution of purified DHX36, with 50 nM as highest concentration was used. In standard treated capillaries (NanoTemper Technologies) microscale thermophoresis analysis was performed with 80% LED, 20% MST power, on the Monollith NT.115 (NanoTemper Technologies). Figures were made with Hill fit using the MO Affinity analysis software.

### 5mer Z-scoring

The BEDTools utility, *getfasta*, was used to recover the genomic nucleotide sequences corresponding to the remaining intervals in these BED files. To produce background, we used an in-house script to scramble the target read sequences while preserving GC content. We then counted the 5mers for target and for background respectively and calculated Z-score enrichment by proportion, 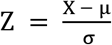. X is the proportion 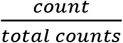 for a given 5mer. Respectively, μ and σ are the average and standard deviation of 5mer proportions in background.

## ACCESSION CODES

The PAR-CLIP, ribosome footprinting, and RNA-seq sequence data have been deposited in the National Center for Biotechnology Information (NCBI) Sequence Read Archive under the accession number GSE105175.

## ACKNOWLEDGMENTS

Research in the Paeschke laboratory is supported by the Emmy-Noether Program of the Deutsche Forschungsgemeinschaft as well as an ERC Stg Grant (638988-G4DSB). Work in the Hafner laboratory was supported by the Intramural Research Program of the National Institute for Arthritis and Musculoskeletal and Skin Disease. We thank the NIAMS Genomics Core Facility and Gustavo Gutierrez-Cruz and Dr. Stefania Dell’Orso (NIAMS/NIH) for sequencing support and Dr. Suman Ghosal (NIAMS/NIH) for binding motif analysis. We thank Dr. Andreas Schlosser for Mass Spectrometry analysis of SILAC experiments and the iPSC/CRISPR Centre (UMCG/ERIBA) for creating the DHX36-KO cell line. We thank Dr. Alexander Buchberger (University of Würzburg) for generous help and support during the whole period of this project and Maria Gallant and Eike Schwindt for experimental support. We thank Dr. Henrike Maatz (MDC Berlin) and Dr. Satyaprakash Pandey (ERIBA, Groningen) for carefully reading of the manuscript.

---

**Supplementary Figure 1:**
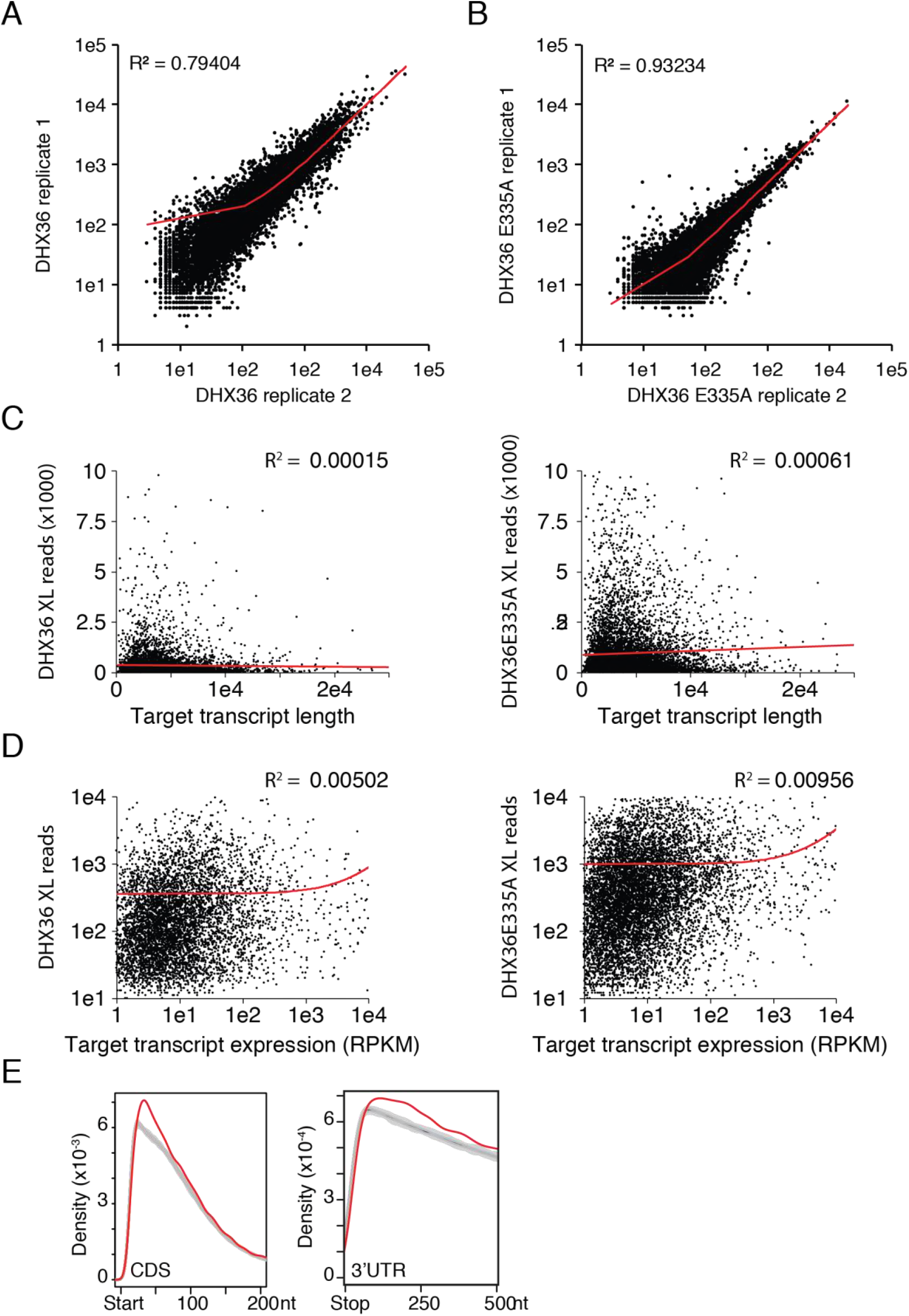
Replicates of DHX36 and DHX36 E335A PAR-CLIP and binding determinants (related to Figure 2). The correlation of crosslinked reads from the biological replicates of DHX36 (**A**) or DHX36 E335A (**B**) PAR-CLIP experiments is shown. **(C)** DHX36 binding on target mRNA is determined neither by target transcript length nor expression. Correlation between crosslinked reads from DHX36 PAR-CLIP (left panels) or DHX36 E335A (right panel) PAR-CLIP and target transcript length. (**D**) Correlation of DHX36 binding with target transcript expression from DHX36 PAR-CLIP (left panels) or DHX36 E335A (right panel). Correlation coefficient (R^2^) are indicated. (**E**) Details of Figure 2E. Metagene analyses of the distribution of DHX36 binding clusters 200 nt downstream of the start codon and 500 nt downstream of the stop codon, respectively. The distribution of 1,000 mismatched randomized controls is shown in grey lines

**Supplementary Figure 2:**
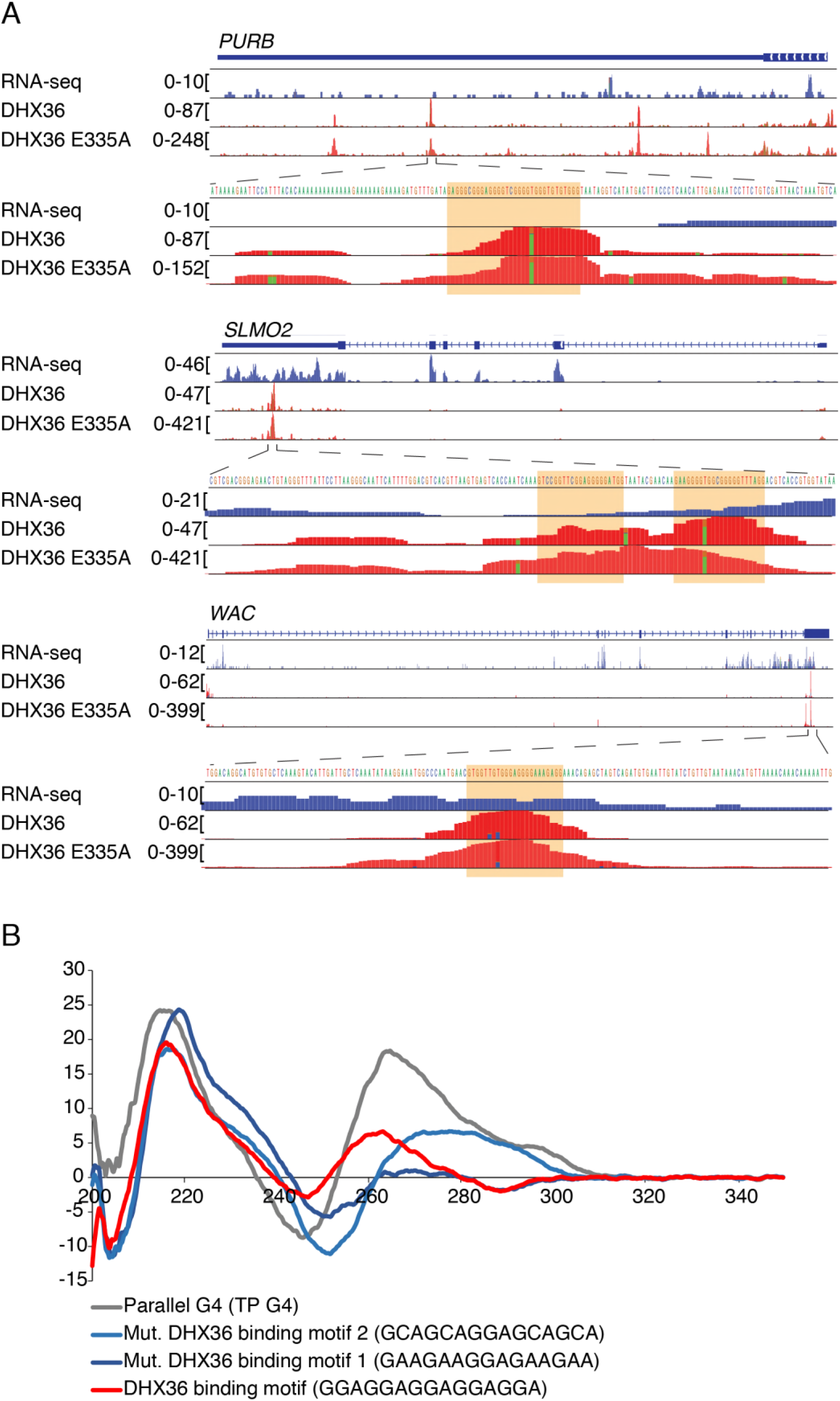
DHX36 binds at G-rich sites and the consensus motif is able to form a parallel G-quadruplex *in vitro* (related to Figure 3). (**A**) Top panels: Screenshots of the DHX36 and DHX36 E335A PAR-CLIP binding sites for the representative target genes PURB, SLMO2, and WAC. The gene structure is shown, as well as coverage from a HEK293 RNA-seq experiment. The bottom two tracks show the alignment of sequence reads with characteristic T-to-C mutations from a DHX36 and DHX36 E335A PAR-CLIP experiment. Bottom panels: Close-up of the indicated 150 nt region in the 3’ UTR of PURB, SLMO2, and WAC. The G-rich DHX36 binding element is highlighted in orange. (**B**) Circular dichroism measurements of oligonucleotides indicated below after performing the G4-folding protocol. The DHX36 PAR-CLIP derived binding motif (red) shifts towards peaks of positive control TP-G4 (grey), whereas a mutated binding motif (light and dark blue) do not shift. Lines represent mean of ten subsequent measurements.

**Supplementary Figure 3:**
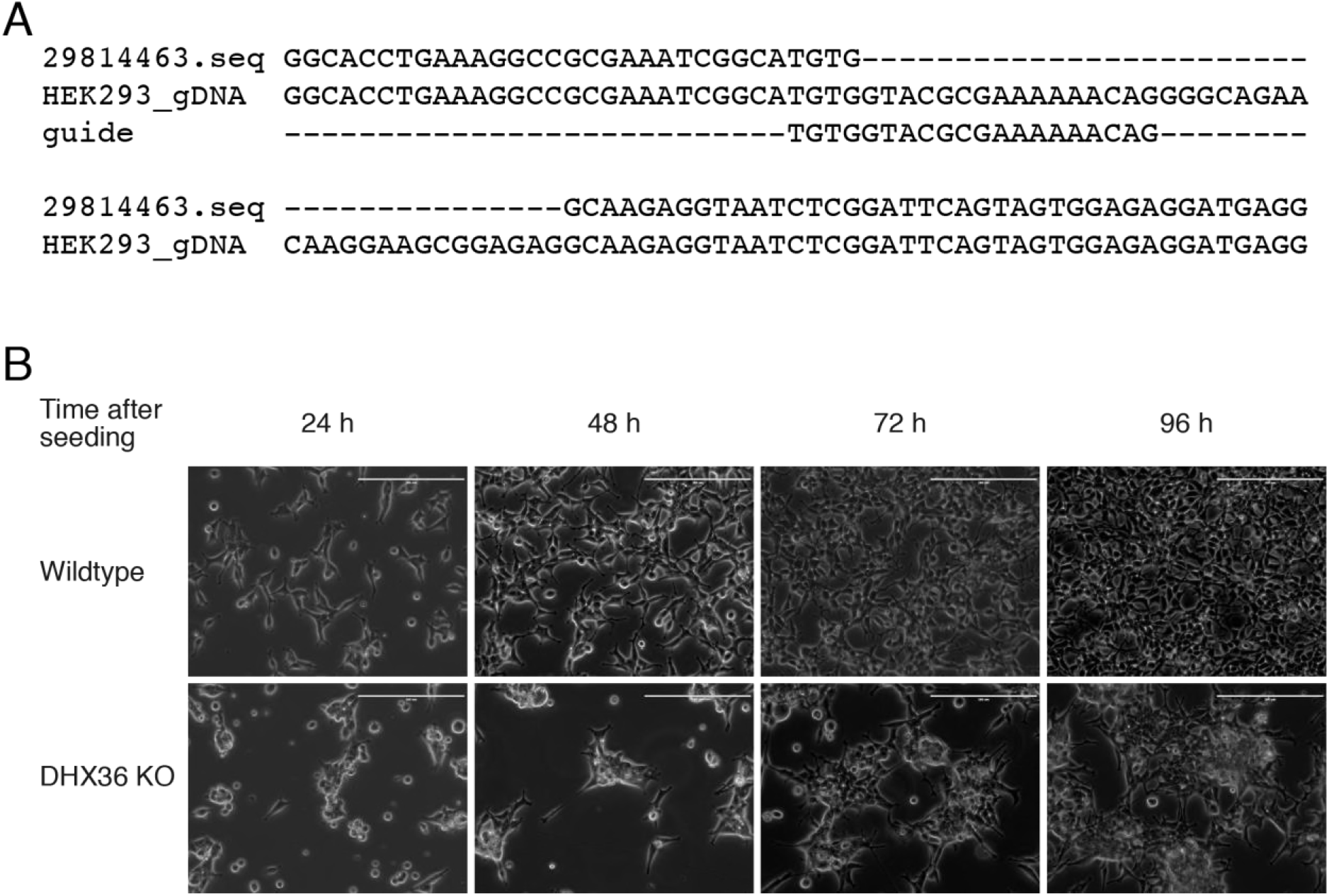
DHX36 KO cell lines (related to Figure 4). **(A)** Sequence of genomic DNA of DHX36 knockout clone indicates the disruption of the gene. Sequence of guide RNA (gRNA) is indicated. **(B)** DHX36 KO cells have a proliferation defect. Changes in morphology of DHX36 knockout cells compared to parental wildtype HEK293 cells. KO cells are not able to equally spread over the culture plate surface. Images were taken at indicate timepoints after seeding. Bar represents nm.

**Supplementary Figure 4:**
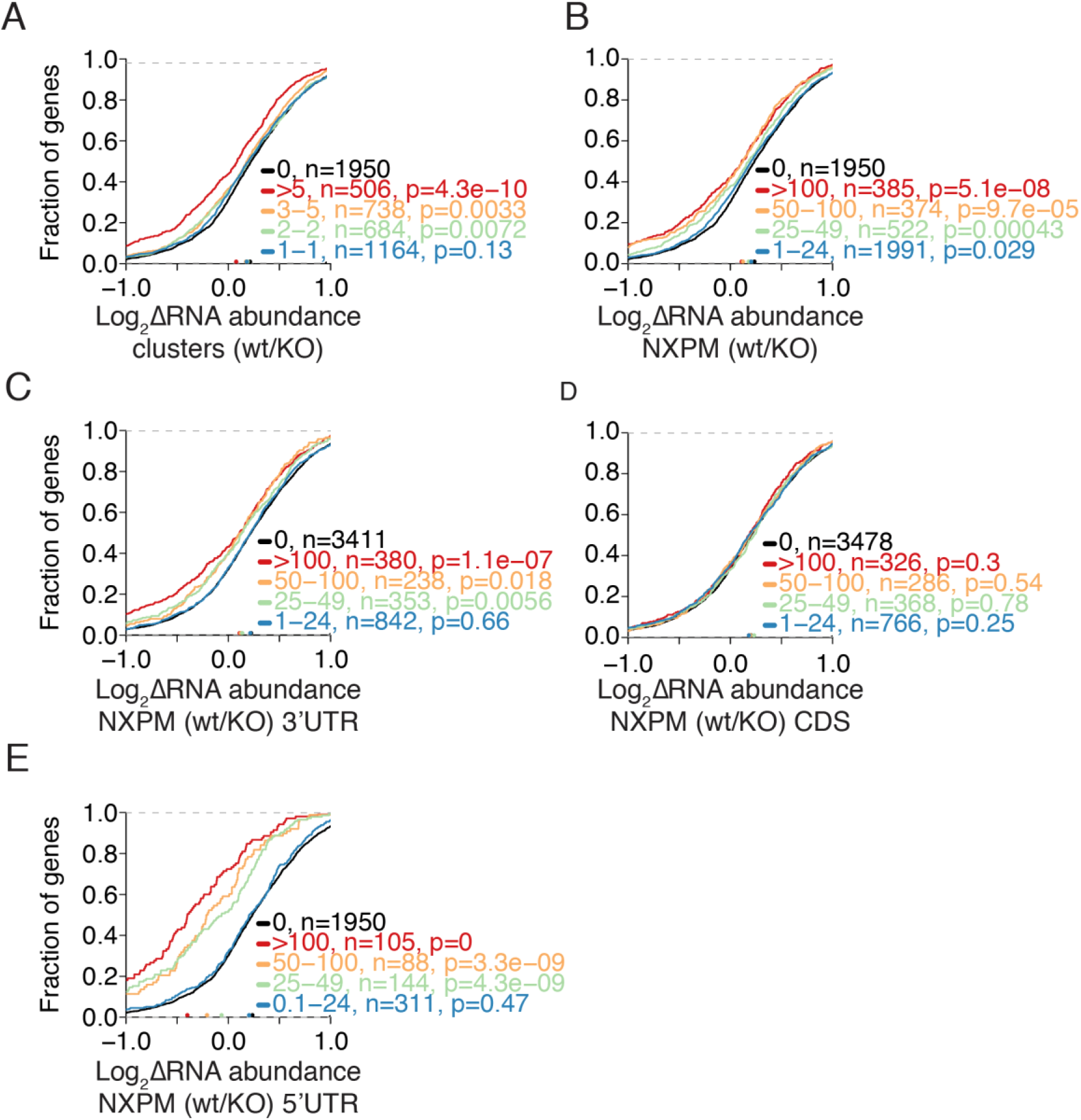
same as Figure 4, but showing data for the wildtype FH-DHX36 (related to Figure 4)

**Supplementary Figure 5 (related to Figure 5):**
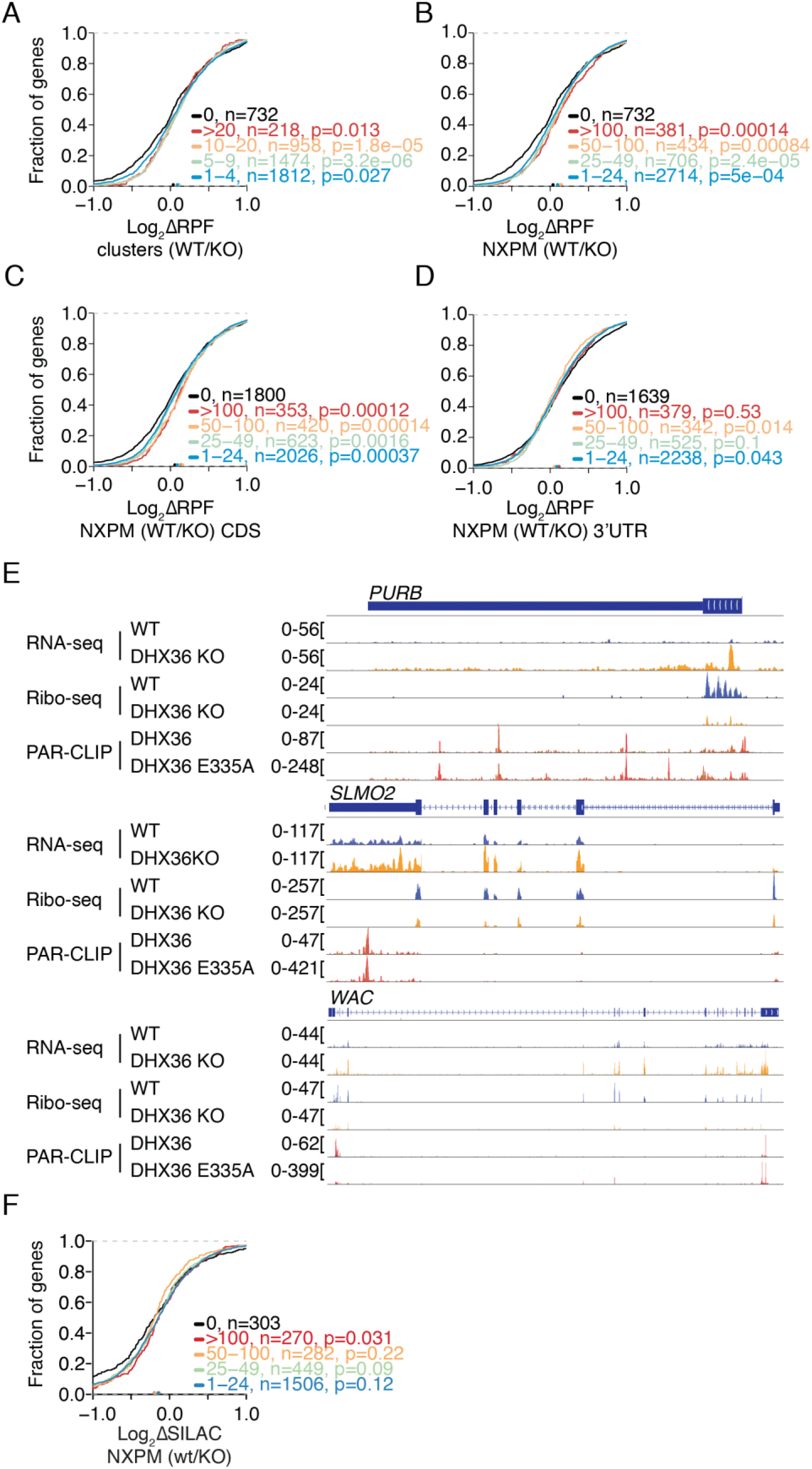
(A-D, F) Same CDFs as in Fig. 5, only with bins representing all DHX36 PAR-CLIP targets, not just top targets. **(E)** Screenshot of RNA-seq and Ribo-seq coverage in wildtype and DHX36 KO HEK293 cells on the representative DHX36 targets PURB, SLMO2, and WAC. Bottom two tracks show the coverage for DHX36 and DHX36 E335A PAR-CLIP.

**Supplementary Figure 6:**
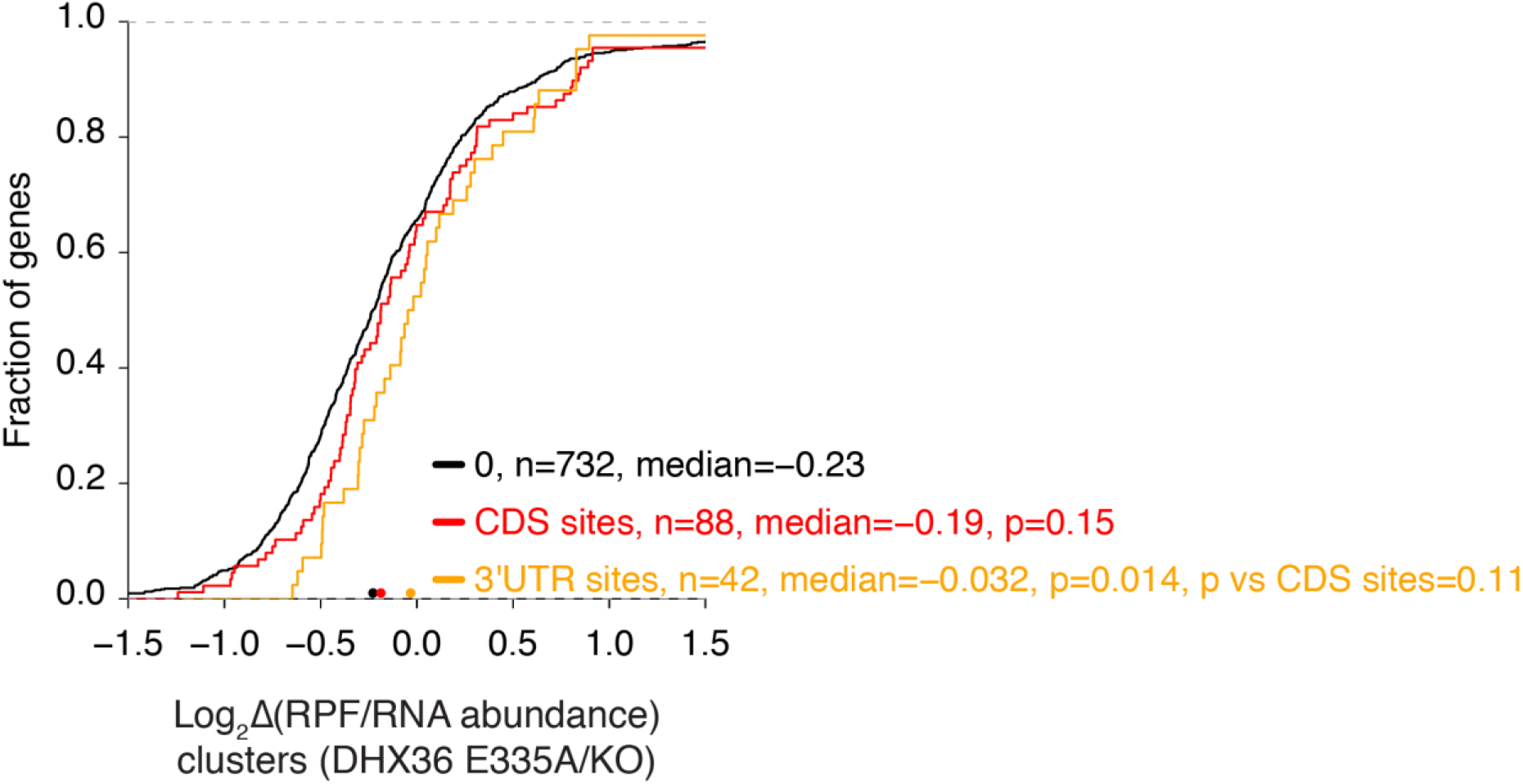
CDS binding by DHX36 does not alter target mRNA abundance (related to Figure 6). Cumulative distribution function comparing changes in DHX36 target mRNA abundance of DHX36 knockout cells (n=3) and parental HEK293 cells (n=3). DHX36-target mRNAs (determined by binding clusters) are separated by coding-sequence bound and 3'UTR bound.

**Supplementary Figure 7:**
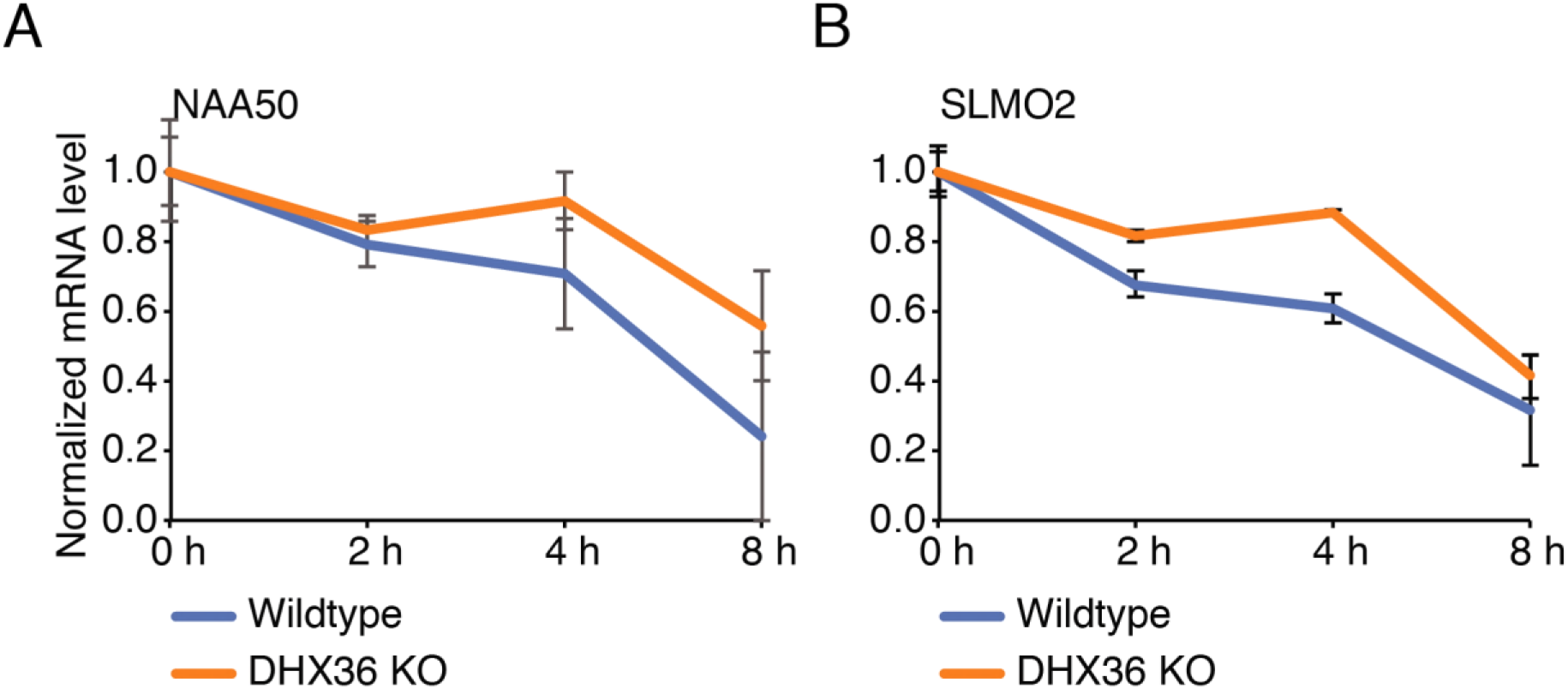
DHX36 regulates target mRNA stability (related to Figure 7) DHX36 target mRNAs increase their half-life upon DHX36 KO shown by qPCR of NAA50 (A) and SLMO2 (B) after transcriptional block using Actinomycin D and isolation of RNA at the indicated timepoints.

**Supplementary Table S1. Overlap of known DHX36 RNA targets with PAR-CLIP clusters and sequencing statistics of PAR-CLIPs. Related to Figure 2.**

**Supplementary Table S2. Occurrence and Z-scores for all 5-mers in the high confidence PAR-CLIP binding sites for DHX36 and DHX36 E335A. Related to Figure 3.**

**Supplementary Table S3. RNA-seq for DHX36 KO and parental HEK293 cells. Related to Figure 4.**

**Supplementary Table S4. Ribo-seq data for DHX36 KO and parental HEK293 cells. Related to Figure 5.**

**Supplementary Table S5. RNA-seq data from chromatin for DHX36 KO and parental HEK293 cells, as well as RNA-seq of total RNA from DHX36 KO and parental HEK293 cells treated with PDS. Related to Figure 7.**

